# *Ataxia-telangiectasia mutated* (*Atm*) disruption sensitizes spatially-directed H3.3K27M/TP53 diffuse midline gliomas to radiation therapy

**DOI:** 10.1101/2023.10.18.562892

**Authors:** Avani Mangoli, Sophie Wu, Harrison Q. Liu, Michael Aksu, Vaibhav Jain, Bronwen E. Foreman, Joshua A. Regal, Loren B. Weidenhammer, Connor E. Stewart, Maria E. Guerra Garcia, Emily Hocke, Karen Abramson, Nerissa T. Williams, Lixia Luo, Katherine Deland, Laura Attardi, Kouki Abe, Rintaro Hashizume, David M. Ashley, Oren J. Becher, David G. Kirsch, Simon G. Gregory, Zachary J. Reitman

**Author notes:** **Lead Author:** Zachary J. Reitman, MD, PhD, 30 Duke Medicine Circle, Box 3085, Durham, 27710, NC, USA. **Corresponding author:** Simon G. Gregory, PhD, 300 N. Duke Street, DUMC 104775, Durham, 27701, NC, USA. **Declaration of Interests Statement:** DGK is a cofounder of and stockholder in XRAD Therapeutics, which is developing radiosensitizers. DGK is a member of the scientific advisory board and owns stock in Lumicell Inc, a company commercializing intraoperative imaging technology. None of these affiliations represent a conflict of interest with respect to the work described in this manuscript. DGK is a coinventor on a patent for a handheld imaging device and is a coinventor on a patent for radiosensitizers. XRAD Therapeutics, Merck, Bristol Myers Squibb, and Varian Medical Systems have provided research support to DGK, but this did not support the research described in this manuscript. ZJR receives royalties for intellectual property related to brain tumor diagnostic tests that is managed by the Duke Office of Licensing and Ventures and has been licensed to Genetron Health, and honoraria for teaching from Oakstone Publishing and Eisai Pharmaceuticals. The other authors have no conflicting financial interests.

## Abstract

Diffuse midline gliomas (DMGs) are lethal brain tumors characterized by p53-inactivating mutations and oncohistone H3.3K27M mutations that rewire the cellular response to genotoxic stress, which presents therapeutic opportunities. We used RCAS/tv-a retroviruses and Cre recombinase to inactivate p53 and induce K27M in the native *H3f3a* allele in a lineage- and spatially-directed manner, yielding primary mouse DMGs. Genetic or pharmacologic disruption of the DNA damage response kinase Ataxia-telangiectasia mutated (ATM) enhanced the efficacy of focal brain irradiation, extending mouse survival. This finding suggests that targeting ATM will enhance the efficacy of radiation therapy for p53-mutant DMG but not p53-wildtype DMG. We used spatial *in situ* transcriptomics and an allelic series of primary murine DMG models with different p53 mutations to identify transactivation-independent p53 activity as a key mediator of such radiosensitivity. These studies deeply profile a genetically faithful and versatile model of a lethal brain tumor to identify resistance mechanisms for a therapeutic strategy currently in clinical trials.

## Introduction

Diffuse midline gliomas (DMGs) are lethal brain tumors of children and young adults. The tumors are localized in essential midline brain structures, such as the brainstem and thalamus, making them surgically inoperable and unresponsive to conventional chemotherapy. The median overall survival of DMGs is less than two years. While radiation therapy may improve symptoms and extend life, it remains palliative. Somatic, activating lysine 27 to methionine mutations in histone variant 3.3 (H3.3K27M) are a defining feature of DMG^1, 2^. Approximately 70% of DMGs harbor inactivating mutations of the tumor suppressor *TP53*^1–3^, which are associated with radioresistance in patients and preclinical models^4, 5^.

A key limitation to current primary DMG preclinical models is the ability to induce K27M mutations in the native *H3f3a* locus in a spatial-, lineage-, and temporally controlled manner. Patient-derived xenografts^6^, patient-derived cell lines^7^, *in utero* electroporation^7^, and syngeneic mouse models^8, 9^ have provided key insights into the disease. A conditional H3f3a-loxP-Stop-loxP-K27M-Tag allele (H3f3a^LSL-K27M-Tag^) has also been generated that allows expression of H3.3K27M from the endogenous mouse *H3f3a* locus in the presence of Cre recombinase^10^. However, this model has been limited by the cell lineages that can be interrogated with existing Cre driver lines, such as *Nestin*-Cre^10^. To date, conditional H3.3K27M alleles have not been interrogated in an entirely spatially controlled manner. We and others have used the RCAS/tv-a retroviral system for spatially-directed modulation of glioma tumorigenesis in the mouse^5, 11–15^. The RCAS/tv-a platform has been used to deliver exogenous H3.3K27M^13, 14, 16^, but to our knowledge it has not been used to edit the endogenous *H3f3a* allele. A variety of these model systems have been used to investigate the mechanisms associated with the development of DMG and assess therapeutic strategies.

Inhibition of the ataxia-telangiectasia mutated kinase (ATM) has emerged as a strategy to enhance the efficacy of radiation therapy for DMG^17^. ATM is a master orchestrator of the DNA damage response to double strand breaks^17^. Patients with hereditary loss-of-function ATM variants, and tumors containing such ATM variants, are exquisitely sensitive to radiation therapy^17^. Consequently, a brain-penetrant ATM inhibitor has entered clinical trials for adult brain tumors (NCT03423628)^18^. A recent study identified ATM inhibition as a potent radiosensitization strategy in a variety of patient-derived pediatric high-grade glioma models^6^. We found that functional ATM loss radiosensitized primary mouse models of DMG driven by p53 loss, but not those that were p53 wildtype^5, 11^. ATM loss increased tumor sensitivity to radiotherapy by radiosensitization of neoplastic cells rather than vasculature^12^. However, it remains uncertain if H3.3K27M affects the ability for *Atm* loss to radiosensitize primary DMG. This is of particular importance since H3.3K27M regulates the important p16 molecular checkpoint that regulates G1-to-S cell cycle progression^13^ and could thereby influence radiation response.

Here, we examine strategies to exploit the genomically-stressed cell state in H3.3K27M/TP53-altered DMG. We improve upon previous models that delivered H3.3K27M from an exogenous RCAS payload^13^ ^11^ by combining the RCAS/tv-a system with H3f3a^LSL-K27M-Tag^ mice to express H3.3K27M from the endogenous *H3f3a* locus. This autochthonous mouse model enabled us to analyze the impact of *Atm* loss in the context of H3.3K27M/TP53-altered brain tumors to mimic human DMG^10^. We found that primary DMGs expressing H3.3K27M and driven by p53 loss were radiosensitized by *Atm* loss. To explore resistance mechanisms in specific tumor cells, we examined primary mouse DMGs after focal brain irradiation using high-resolution single cell *in situ* sequencing (ISS). The result identified overexpression of the cell cycle regulator *Cdkn1a* as a putative resistance factor in *Atm*-intact DMG. We showed that *Cdkn1a*, or transcriptional activity of p53 in general, is dispensable for DMG radiosensitization by *Atm* loss. Therefore, non-transactivation functions of p53 may determine sensitivity of DMGs to combinations of ATM inhibitors and radiation therapy. The high-resolution results describe a genetically faithful and flexible primary mouse model of DMG, identifying mechanisms of resistance to a therapeutic strategy currently in clinical trials.

## Results

### Conditional p53 loss and H3.3K27M expression in retrovirus-induced mouse DMGs

To express H3.3K27M from the endogenous *H3f3a* locus in retrovirus-induced primary mouse gliomas, we used a H3f3a^LSL-K27M-Tag^ allele that expresses H3.3K27M in the presence of Cre recombinase^10^. To incorporate the H3f3a^LSL-K27M-Tag^ allele into the RCAS/tv-a retrovirus system, mice were bred with Nestin^TVA^ mice to allow RCAS retroviruses to specifically transduce TVA+ *Nestin-*expressing neural stem cells. To investigate deletion of p53 specific to tumors, we crossbred a p53 variant in which critical exons were flanked by loxP sites (floxed or FL) allowing for the functional deletion of p53 in the presence of Cre recombinase. We first introduced retroviruses into Nestin^TVA^; p53^FL/FL^; H3f3a^LSL-K27M-Tag/+^ mice and compared these to matched mice lacking the H3f3a^LSL-K27M-Tag^ allele (Figure 1A). We induced DMGs by injecting mice with RCAS retroviruses expressing Cre recombinase, firefly luciferase, the oncogene platelet-derived growth factor ligand beta (PDGF-B) and monitored for tumor formation via *in vivo* imaging (Figure 1B). Using luciferase-based bioluminescent imaging to detect tumors, we determined there was no difference in time to tumor formation in H3f3a^LSL-K27M-Tag/+^ mice compared to matched mice lacking the H3f3a^LSL-K27M-Tag^ allele (Figure 1C). Tumors exhibiting hypercellularity and diffuse infiltration of nearby normal brain on H&E formed within 4-8 weeks with high penetrance (Figure 1D). We detected HA expression indicating the presence of the HA tag that is present on both H3.3K27M and PDGF-B constructs (Figure 1E). As expected, p53 was not detected in these p53^FL/FL^ tumors by IHC (Figure 1F). Histone 3 lysine 27 trimethylation was dramatically decreased in the H3f3a^LSL-K27M-Tag/+^ tumors compared to controls (Figure 1G), indicating that H3.3K27M could functionally deplete H3K27me3 as predicted^19^. Ki67 was elevated to >50% of tumor cells regardless of H3.3K27M status (Figure 1H). Anti-Flag immunohistochemistry confirmed the presence of the Flag tag on the H3.3K27M construct (Figure 1I). Flag IHC demonstrated H3.3K27M-Tag+ cells diffusely infiltrating from a hypercellular tumor core into brain parenchyma (Figure 1J), suggesting diffuse, infiltrative biology seen in human DMG. These results demonstrate that RCAS/tv-a and a conditional H3f3a^LSL-K27M-Tag^ allele can be combined to target K27M to the *H3f3a* gene in time, lineage, and space to generate primary mouse DMGs.

**Figure 1.**
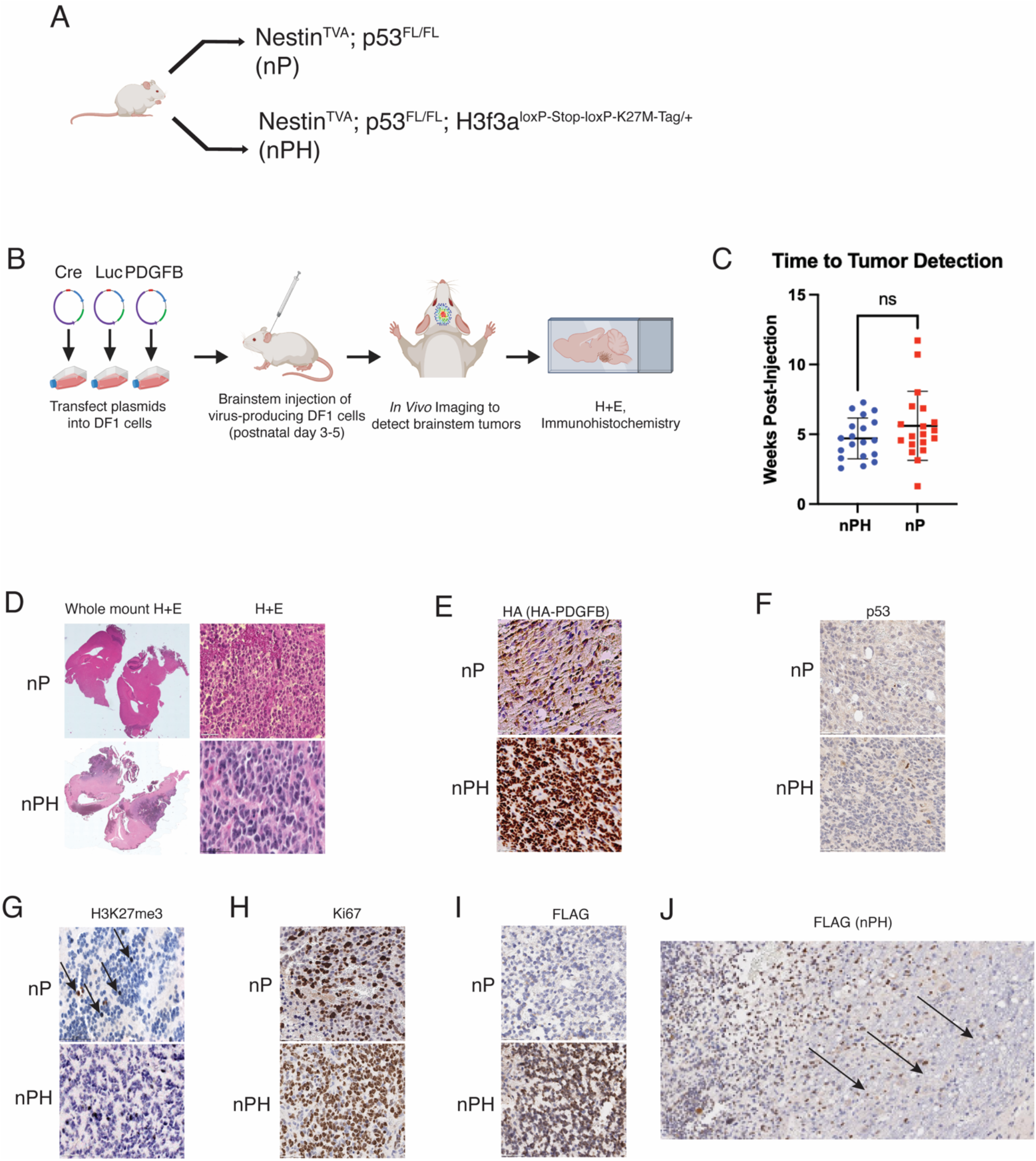
Primary murine diffuse midline gliomas generated using a conditional H3.3K27M allele and RCAS/tv-a retroviruses system. (A) Representation of mouse genotypes used to generate primary mouse DMGs with p53 loss (Nestin^TVA^; p53^FL/FL^) and mouse DMGs with p53 loss and H3.3K27M (Nestin^TVA^; p53^FL/FL^; H3f3a^loxP-Stop-loxP-K27M-Tag/+,^ nPH) with or without conditional H3.3K27M allele. Mice also contained one intact and one floxed allele of Atm (Atm^FL/+^, not shown). (B) Procedure for intracranial injection of RCAS viruses, followed by tumor detection and tissue-based studies of mice bearing tumors. (C) Dot plot showing time to tumor formation between nPH and nP mice without any statistical significance. Statistical test utilized Mann-Whitney test. (D) Whole mount and H&E slides showing tumor within nP and nPH mice. (E) IHC for HA expression indicating the presence of the PDGF-B HA tag. (F) IHC for p53. (G) IHC for histone 3 lysine 27 trimethylation (H3K27me3). (H) IHC displaying the Ki67 proliferation of tumors. (I) Anti-Flag IHC confirmed the presence of the Flag-HA tag. (J) Anti-Flag IHC at tumor periphery shows infiltration of tumor cells into normal brain parenchyma identified by arrows. Scale bar for all H&E and IHC images (C-H) = 50 µm. Scale bar for Whole Mount = 200 uM. Scale bar for FLAG IHC infiltrating biology (I) = 500 µm

### *Atm* loss radiosensitizes primary p53-null/H3.3K27M DMGs

Targeting the ATM kinase has emerged as a potential strategy to increase the efficacy of standard-of-care radiation therapy for brain tumors^5, 6, 17^. We sought to determine if disruption of ATM could radiosensitize primary mouse DMGs containing p53 and H3.3K27M alterations. Previously, we established that *H3f3a*-wildtype brainstem gliomas lacking *Atm* in the tumor cells were radiosensitized compared to littermate controls with a functional *Atm* allele in their tumors^5^. However, these mice lacked H3.3K27M which disrupts the G1-to-S cell cycle checkpoint^13^ and may thereby affect the downstream effects of ATM deficiency^17^. We hypothesized that *Atm* inactivation in the presence of the H3.3K27M allele would also radiosensitize tumors. To test this, we examined tumor-free survival of Nestin^TVA^; p53^FL/FL^; H3f3a^LSL-K27M-Tag/+^; Atm^FL/FL^ (nPHA^FL/FL^) mice and compared these to controls with intact ATM in their tumors of genotype Nestin^TVA^; p53^FL/FL^; H3f3a^LSL-K27M-Tag/+^; Atm^FL/+^ (nPHA^FL/+^) (Figure 2A). There was no difference in tumor-free survival between nPHA^FL/FL^ and nPHA^FL/+^ mice in the absence of irradiation (Figure 2B). To test if *Atm* deletion radiosensitizes p53-null/H3.3K27M DMGs, we delivered three daily fractions of 10 Gy focal brain irradiation to mice using a Small Animal Radiation Research Platform (SARRP). nPHA^FL/FL^ mice had significantly longer median survival compared to nPHA^FL/+^ mice (P-value=0.03 using Mantel Cox (log rank test), Figure 2C). Thus, *Atm* deletion in tumor cells enhanced the efficacy of focal brain irradiation for primary p53-null/H3.3K27M DMGs. IHC confirmed HA expression (Figure 2D, E), p53 loss (Figure 2F), H3.3K27M presence (Figure 2G), Ki67 (Figure 2H) and Anti-FLAG (Figure 2I). These results show that *Atm* disruption enhances the efficacy of radiation therapy for primary mouse DMGs that contain p53 loss and H3.3K27M mutation.

**Figure 2.**
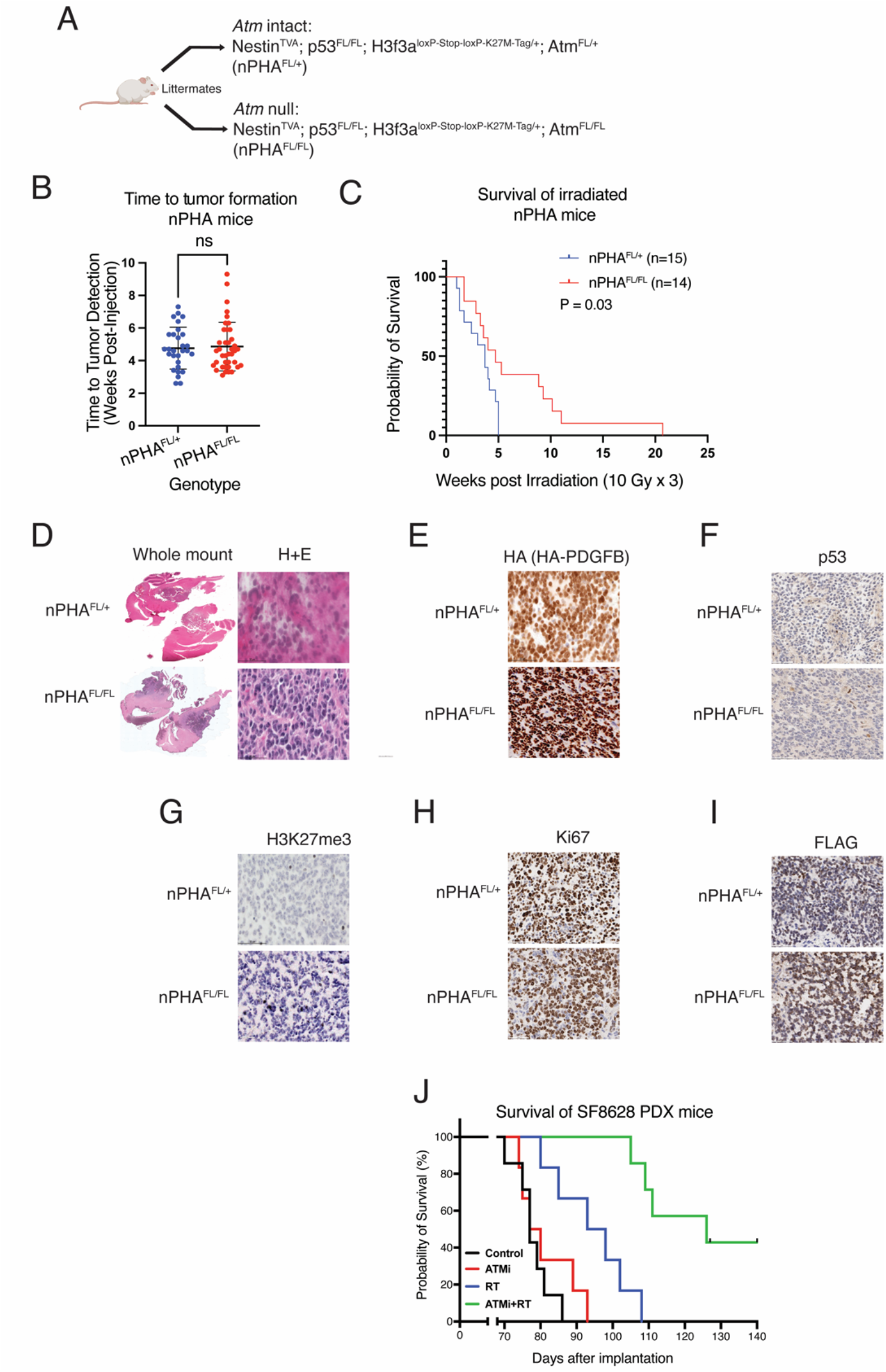
Atm loss radiosensitizes primary H3.3K27M/TP53 mouse DMGs. (A) Schematic showing nPHA^FL/+^ (Atm^FL/+^) and nPHA^FL/FL^ (Atm^FL/FL^) within RCAS/TVA retrovirus and conditional H3K27M allele. (B) Time to tumor formation showing no statistical difference between nPHA^FL/FL^ and nPHA^FL/+^ mice. Statistical test utilized Welch’s t-test. (C) Overall survival of between nPHA^FL/FL^ and nPHA^FL/+^ mice following three daily fractions of 10 Gy image-guided focal brain irradiation shows significantly longer median survival in nPHA^FL/FL^ (p-value = 0.03) using Mantel Cox (log rank test). (D) Whole mount and H&E slides showing tumor within (top) nPHA ^FL/+^ and (bottom) nPHA^FL/FL^ mice exhibiting hypercellularity and infiltration of normal brain. (E) IHC for HA expression indicating the presence of the PDGF-B HA tag. (F) IHC showing loss of p53 in nPHA ^FL/+^ and nPHA^FL/FL^ mice (G) IHC showing histone 3 lysine 27 trimethylation (H3K27me3). (H) IHC displaying the Ki67 proliferation of tumors. (I) Anti-Flag IHC confirmed the presence of the Flag-HA tag. (J) Overall survival of mice bearing SF8628 diffuse midline glioma patient-derived xenografts were treated with 20mg/kg of AZD1390 for 2 weeks (ATMi, 5 days a week x 2 weeks) and/or focal brain irradiation (RT, 2 Gy x 3 days a week for 12 Gy total). Scale bar for all H&E and IHC images = 50 µm

To validate these findings, we confirmed that pharmacologic inhibition of ATM could radiosensitize human patient-derived model of DMG. To do so, we tested if combination of the brain-penetrant ATM inhibitor AZD1390^18^ combined with focal brain irradiation could similarly improve survival of a patient-derived xenograft model of H3.3K27M-mutant and p53-mutant diffuse midline glioma, SF8628^20–23^. The combination of AZD1390 and irradiation significantly extended median survival compared to either treatment alone (Figure 2J). These results confirm that either pharmacologic or genetic targeting of ATM can radiosensitize multiple types of *in vivo* DMG models.

### In situ multiplexed microscopy reveals cell cycle and Semaphorin pathway changes after irradiation and *Atm* disruption

To explore mechanisms of radiation efficacy and resistance, we performed spatially resolved gene expression analyses of primary mouse DMGs. Our previous work identified key differences in response to irradiation and to *Atm* loss between the neoplastic and vascular compartments within primary mouse tumors^12^. To distinguish compartment-specific changes in gene expression, such as vascular and immune cells in specific regions of tumor and nontumor brain, we needed to profile expression changes at single-cell resolution and in a spatially resolved manner. To achieve such resolution, we used 10xGenomics Xenium ISS platform to profile primary p53-null/H3.3K27M mouse DMGs. We examined DMG-bearing mice treated with or without focal brain irradiation (10 Gy x 3), and with or without tumoral *Atm* loss as depicted in Figure 3A. We examined 5 μm mid-sagittal sections of formalin-fixed, paraffin-embedded (FFPE) tumor-bearing brains. We supplemented 10xGenomics’ standard mouse brain content with a custom panel containing padlock probes resulting in 298 brain- and DMG-specific mRNA transcript assays (Table S1) which were hybridized, amplified, and then *in situ* sequenced within each sample. Individual cells were detected by nuclear DAPI staining and cell boundaries were segmented by expanding outwards until 15 μm or the boundary of another cell was reached (see Methods). This yielded 790,374 individual cells across four tumor-bearing brains. Next, we clustered cells based on their transcriptional profiles and compared cell type composition between samples. Uniform Manifold Approximation and Projection (UMAP)^24^ reduction and projected differentiated normal and neoplastic brain cells into 20 and 29 clusters per specimen, respectively (Figure S1). Examination of differentially expressed marker genes in each cluster identified neoplastic and normal cells including GABAergic interneurons marked by *Gad1* and *Gad2*; microglia marked by *P2ry12*, *Lyz2* and *C1qa*; and endothelial cells marked by *Cd34, Fn1*, and *Adgrl4* (Figure S2). We used canonical cell type markers and label transfer-based methods to collapse cell clusters into 10 cell archetypes (neoplastic, endothelial, neuron, astrocyte, oligodendrocyte, microglia, T-lymphocyte, etc.) that could be directly compared across specimens (Table S2 and Figure S3). This analysis revealed masslike tumors with infiltrating edges, recapitulating diffuse glioma biology (Figure 3B). Of note, an *Atm*-null post-irradiation tumor was smaller and involuted, which is suggestive of rapid treatment response.

**Figure 3.**
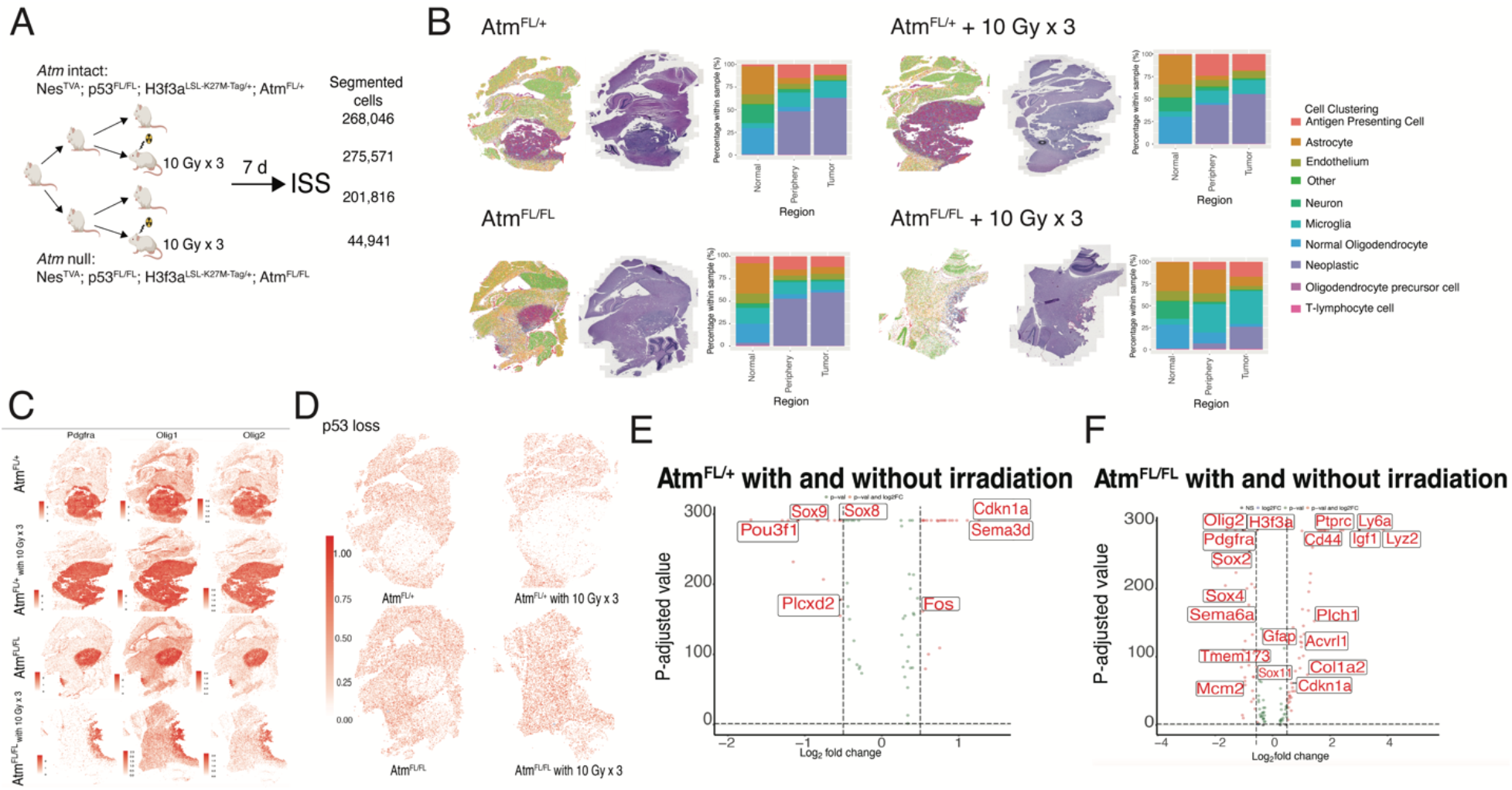
Spatial clustering and differentially expressed genes in primary mouse DMGs treated with focal brain irradiation or tumoral *Atm* deletion. (A) Schematic of DMG-bearing mice with *Atm* intact or null within the tumors, with and without focal brain irradiation that underwent in situ spatial transcriptomic sequencing (ISS). All mice were Nestin^TVA^; p53^FL/FL^; H3f3a^loxP-Stop-loxP-K27M-Tag/+^ with either *Atm* intact (Atm^FL/+^) or *Atm* null (Atm^FL/FL^) in the tumor. (B) Spatial clustering of cells into n=10 cell archetypes based on label transfer in n=4 tumor-bearing mouse brains (*left*), H&E of whole brain (*middle*), distribution of cells within normal brain, tumor periphery and tumor core annotated in bar graph (*right*). Top row indicates *Atm* intact with and without irradiation. Bottom row indicates *Atm* null with and without irradiation. Color legend to the right corresponds to individual cell type noted on bar graph. (C) Spatial identification of tumors by expression of *Pdgfra*, *Olig1*, and *Olig2* within all conditions: *Atm* intact, *Atm* intact with irradiation, *Atm* null, *Atm* null with irradiation (*top to bottom)*. (D) Spatial identification of p53 loss in all tumor conditions: *Atm* intact without and with irradiation (*top row, left to right)*. *Atm* null without and with irradiation (*bottom row, left to right)*. (E) Key differentially expressed genes in *Atm*-intact neoplastic tumor cells treated with and without focal brain irradiation. Log_2_ fold change and p-value for all genes in Table S3. (F) Key differentially expressed genes in *Atm*-null neoplastic tumor cells treated with and without focal brain irradiation. Log_2_ fold change and p-value for all genes in Table S4. All cells within entire slide were used for distribution of cells identified in Figure 3b. Tumor core and periphery were utilized to identify the key differentially expressed genes (Figure E-F)

We used spatially resolved expression data to identify differentially expressed genes among neoplastic cells within the tumor cores. We first localized the tumor core within the full-brain sagittal sections using canonical DMG neoplastic cell markers, *Olig1*, *Olig2*, and *Pdgfra* (Figure 3C). As expected, we could not detect *p53* in the neoplastic cells within tumor core in the Tp53^FL/FL^ model, while low baseline levels could be detected in non-neoplastic cell types (Figure 3D). Similarly, *Atm* transcripts were nearly undetectable in neoplastic cells from the *Atm*-null tumor (mean fold-change −0.636, P<0.0001 when compared to *Atm*-intact tumor). The tumor core, periphery, and nontumor areas were contoured using these data to allow comparisons between matching cell types and locations after irradiation or *Atm* loss (Figure 3B). To identify transcripts that may be differentially expressed after irradiation and/or *Atm* loss, we interrogated differentially expressed genes in neoplastic cells after focal brain irradiation within *Atm* intact tumors (Table S3) *and Atm*-null tumors (Table S4). *Cyclin-dependent kinase 1a (Cdkn1a)*, which encodes p21 a potent regulator of cell cycle progression at G1, was the most differentially expressed gene after focal brain irradiation among *Atm* intact tumors (log-fold change 0.8, P-value = 0 by Wilcoxon test, Figure 3E). *Cdkn1a* was still upregulated, albeit to a lesser degree, after focal brain irradiation among *Atm*-null tumors (log-fold change 0.6, P-value = 5.46E-08 by Wilcoxon test, Figure 3F). Conversely, transcription factors associated with developmental cell states such as *Sox8* and *Sox9* were significantly downregulated after irradiation in the *Atm*-intact tumors, while *Sox*2, *Sox4*, *Pdgfra*, and *Olig2* associated with early glial differentiation were all significantly downregulated after irradiation in the *Atm*-null tumors. These results identify differential expression of cell cycle regulators and of cell-fate-regulating transcription factors after irradiation in a primary DMG mouse model.

Irradiation and *Atm* loss also associated with changes in the expression of Semaphorin genes, which encode secreted and membrane proteins that play critical roles in neuronal wiring and glioma biology. Specifically, *Semaphorin 6A* (*Sema6a*) and *Semaphorin 3D* (*Sema3d)* have been implicated in proliferation and survival of glioma mouse models and glioblastoma^25, 26^. After irradiation in *Atm*-intact tumors, *Sema3d* was significantly increased (log-fold change 1.13, P-value = 0) suggesting that radiation therapy may influence proliferation within the neoplastic core. After irradiation in *Atm*-null tumors, *Sema6a* was significantly decreased (log-fold change −0.40, P-value = 6.59E-15). These results indicate that primary DMG mouse models recapitulate Semaphorin-related molecular pathways in glioma, and that expression of specific Semaphorin genes is specifically altered in neoplastic cells after radiotherapy.

### Neighborhood analysis shows altered immune-neoplastic interactions after treatment

We next examined whether proximity between neoplastic cells and normal cells varied across irradiated or *Atm*-null tumors. Targeting ATM combined with irradiation can bridge innate immune processes with the adaptive immune processes in extracranial cancers^27, 28^. This led us to hypothesize that co-localization of neoplastic cells with microglia and with other antigen-presenting cells would be increased in irradiated and/or *Atm*-null tumor-bearing brains. The single-cell resolution of the ISS data allowed us to carry out neighborhood analysis to quantify spatial proximity between different cell types (Figure 4A-D). Spatially resolved data was used to estimate the mean distance between neoplastic cells and other cell types (Figure 4E-G). Neighborhood analysis identified increased proximity of neoplastic cells and immune cells, such as antigen presenting cells and microglia, after *Atm* loss and after treatment with irradiation, which was especially pronounced in the irradiated *Atm*-null tumor (Figure 4H). Co-localization analysis between neoplastic cells and other cell types confirmed that microglia and antigen-presenting cells were most enriched within 0-500 μm (Figure 4I-K), and again these cell types were most co-localized in the irradiated, *Atm-*null tumor (Figure 4L). These findings suggest that *Atm* loss with irradiation affects the spatial relationship between neoplastic cells the immune microenvironment.

**Figure 4.**
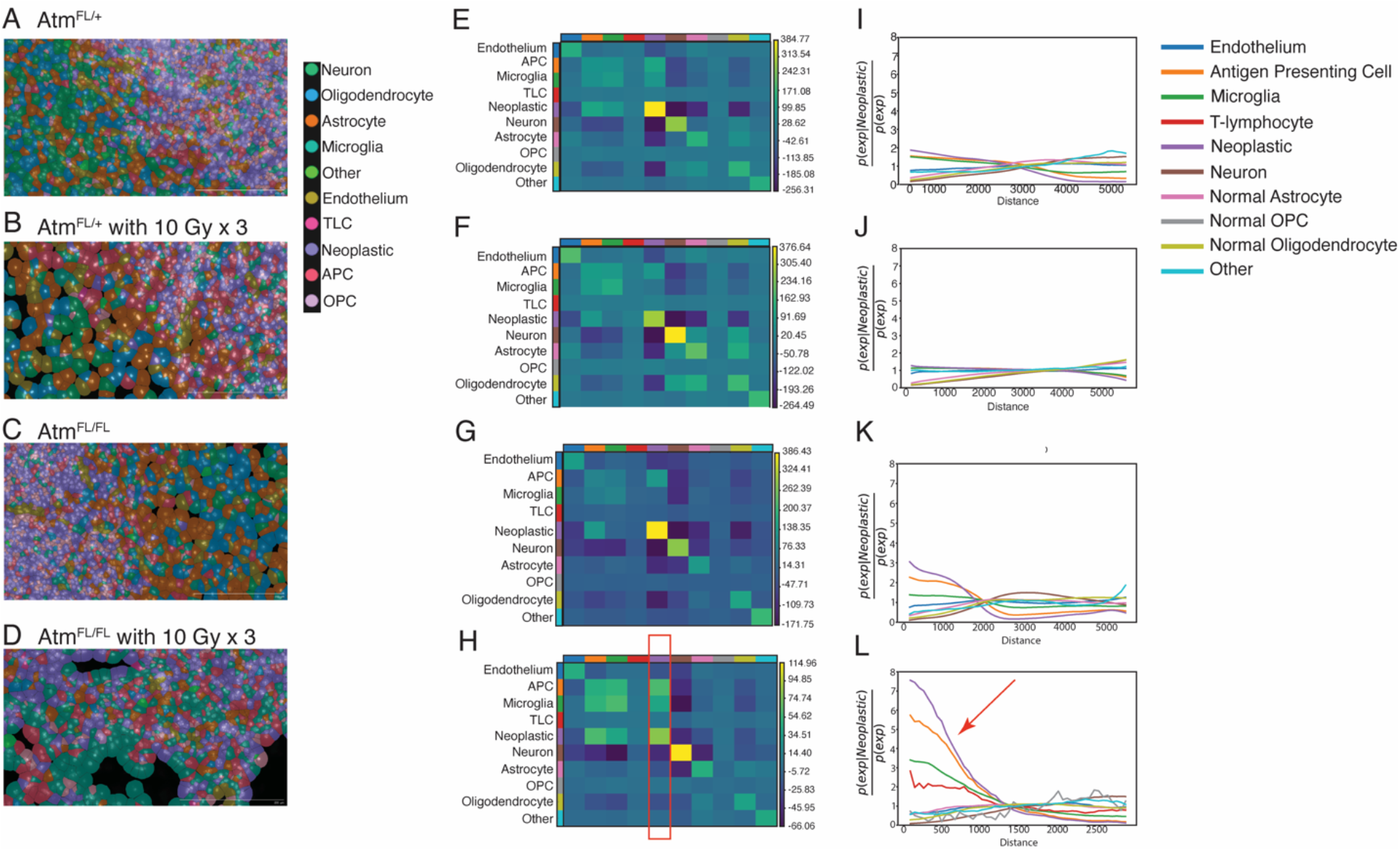
Neighborhood analysis of primary mouse DMGs with tumoral *Atm* loss and/or focal irradiation. (A) Representative image of *Atm*-intact (nPHA^FL/+^) tumor highlighting individual cell types identified by ISS at the border of normal brain and tumor. (B) Representative image of *Atm*-intact (nPHA^FL/+^) tumor with irradiation highlighting individual cell types at the border of normal brain and tumor. (C) Representative image of *Atm*-null (nPHA^FL/FL^) tumor highlighting individual cell types. (D) Representative image of *Atm*-null (nPHA^FL/FL^) tumor with irradiation (E) Neighborhood enrichment analysis of Atm-intact (nPHA ^FL/+^) tumor showing proximity of various cell types in relationship to neoplastic cells. (F) Neighborhood enrichment analysis of *Atm*-intact (nPHA ^FL/+^) tumor with irradiation. (G) Neighborhood enrichment analysis of *Atm*-null (nPHA^FL/FL^) tumor. (H) Neighborhood enrichment analysis of *Atm*-null (nPHA^FL/FL^) tumor with irradiation showing proximity of various cell types in relationship to neoplastic cells. Red box indicates presence of antigen-presenting cells and microglia in relationship to neoplastic cells. (I) Co-occurrence plot of *Atm*-intact (nPHA ^FL/+^) tumor showing number compared to distance of various cell types in relation to neoplastic cells. (J) Co-occurrence plot of *Atm*-intact (nPHA ^FL/+^) tumor with irradiation showing number compared to distance of various cell types in relation to neoplastic cells. (K) Co-occurrence plot of *Atm*-null (nPHA^FL/FL^) tumor showing number compared to distance of various cell types in relation to neoplastic cells. (L) Co-occurrence plot of *Atm*-null (nPHA^FL/FL^) tumor with irradiation showing number compared to distance of various cell types in relation to neoplastic cells. Red arrow indicates increased frequency of immune cells compared to neoplastic cells. Color Legend for Figure (A-D) on left side panel. Color Legend for Figure (E-L) on right side panel. APC – Antigen Presenting Cell. OPC – Oligodendrocyte precursor cell. TLC-T Lymphocyte cell. Neighborhood enrichment and co-occurrence analysis were conducted on entire slide. All unlabeled cells were removed for analysis. Figure (A-D) Scale bar = 200 uM.

### Ligand-receptor analysis reveals endothelial cell communications

We next interrogated cell:cell and cell:ligand receptor interactions in primary mouse DMGs (Figure 5 and Table S5). In all tumor conditions, the endothelial cells had the highest frequency of interactions (Figure 5A-D). We evaluated for statistically significant ligand-receptor interactions (p-value <0.05) amongst the tumors. We noticed that the interaction between endothelium, microglia, and neoplastic cells with Col1a2:CD93 receptor decreased after *Atm* loss and irradiation (Figure 5E-H). CD93 plays a role in tumor-associated vasculature^29^ and changes in expression of Col1a2 has been observed post radiotherapy in other cancers^30^. These provide insight into changes in endothelial cell interactions after tumor irradiation. After irradiation of *Atm*-intact tumors, the cell:ligand interaction of Sema3a:NRP2 between neoplastic cell and microglia decreases (Figure 5I-J). This interaction has been noted to have effects on glioma cell migration^31^ implying a potential alteration in migration with irradiation. The opposite effect is noted in *Atm*-null tumors after irradiation (Figure 5K-L). Thus, ligand-receptor analysis of ISS data suggests that glioma-linked collagen and Semaphorin interactions can be examined in primary DMG mouse models.

**Figure 5.**
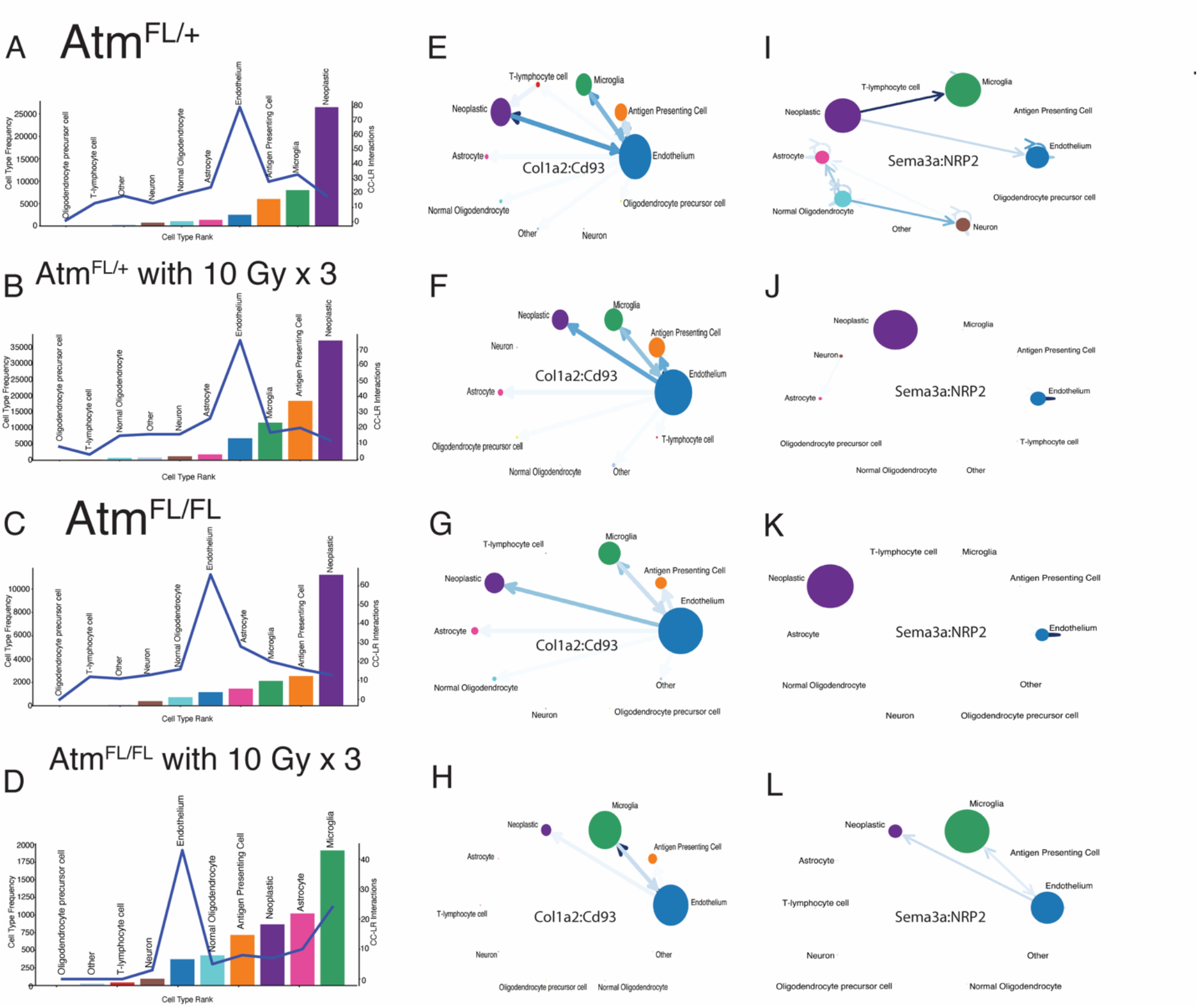
Spatial cell interactions within tumor environment of primary mouse DMGs with tumoral *Atm* loss and/or focal irradiation. (A-D) Bar graph demonstrating number of interactions with cells and ligand receptors of each cell type when compared to frequency of cells. (E-H) Col1a2:Cd93 interaction amongst all cell types between *Atm* null (nPHA^FL/FL^) and *Atm* intact (nPHA^FL/+^). P-value < 0.05. (I-L) Sema3a:NRP2 interaction amongst all cell types between neoplastic cells and endothelium between *Atm* null (nPHA^FL/FL^) and *Atm* intact (nPHA^FL/+^). P-value < 0.05. Cell:Cell and Cell:Ligand interactions were conducted on tumor core and periphery. All unlabeled cells were removed for analysis.

### *Atm* radiosensitizes *Cdkn1a*-null primary murine DMGs

We next dissected specific functions of p53 that may affect the radiosensitivity of mouse DMG. Our primary models of DMG indicated that the presence of functional p53 is a key determinant of whether the tumors are radiosensitized by *Atm* loss. For instance, primary p53-null/H3.3K27M tumors and primary p53-null/*H3f3a*-wildtype tumors are radiosensitized by *Atm* loss (Figure 2C and Deland *et al.* ^5^, respectively). Conversely, p53 wildtype primary DMG models driven by Ink4A/ARF loss or by PTEN loss are not radiosensitized by *Atm* loss^5, 11^. However, it is unknown whether loss of p53 transcriptional activation and/or loss of other p53 functions enables radiosensitization by *Atm* loss. Our ISS data identified increased *Cdkn1a* expression after radiation in neoplastic cells. Since *Cdkn1a* (encoding p21) is a major transcriptional target of p53^32^, we hypothesized that loss of *Cdkn1a* function downstream of p53 may be a key determinant of whether *Atm* loss can radiosensitize primary DMGs. To test if *Cdkn1a* loss allows primary mouse DMGs to be radiosensitized by *Atm* loss, we examined our model of p53-wildtype DMGs driven by Ink4A/ARF loss, which are not radiosensitized by *Atm* loss (Nestin^TVA^; Ink4A/ARF^FL/FL^)^5^. To test if p21 loss could radiosensitize these mice when *Atm* is lost, we bred mice with constitutive p21 loss into this genotype (Nestin^TVA^; p21^-/-;^ Ink4A/ARF^FL/FL^; Atm^FL/FL^ (nIp21A^FL/FL^). We tested the effects of tumor-specific *Atm* loss by comparing these mice to littermate controls with intact *Atm* with genotype Nestin^TVA^; p21^-/-;^; Ink4A/ARF^FL/FL^; Atm^FL/FL^ (nIp21A^FL/+^) (Figure 6A). Time to tumor formation was similar regardless of the presence of intact *Atm* (Figure 6B). Surprisingly, p21-null mice bearing tumors with *Atm* deletion had a shorter survival following fractionated focal brain irradiation compared to littermate controls with intact *Atm* in the tumors (Figure 6C, P<0.03, log-rank test). We confirmed p21 loss by IHC (Figure 6D-E). p21-null tumors with and without *Atm* loss had similar proliferation indices as assessed by Ki67 staining (Figure 6F). TUNEL staining of irradiated tumors showed that *Atm* loss was associated with significantly increased TUNEL staining (P < 0.05, Figure 6G,H), suggesting that tumors lacking both *Atm* and *Cdkn1a* may be primed for apoptosis. These results show that functional *Cdkn1a* is not the key mediator of radiosensitization by *Atm* loss in primary mouse model of DMG.

**Figure 6.**
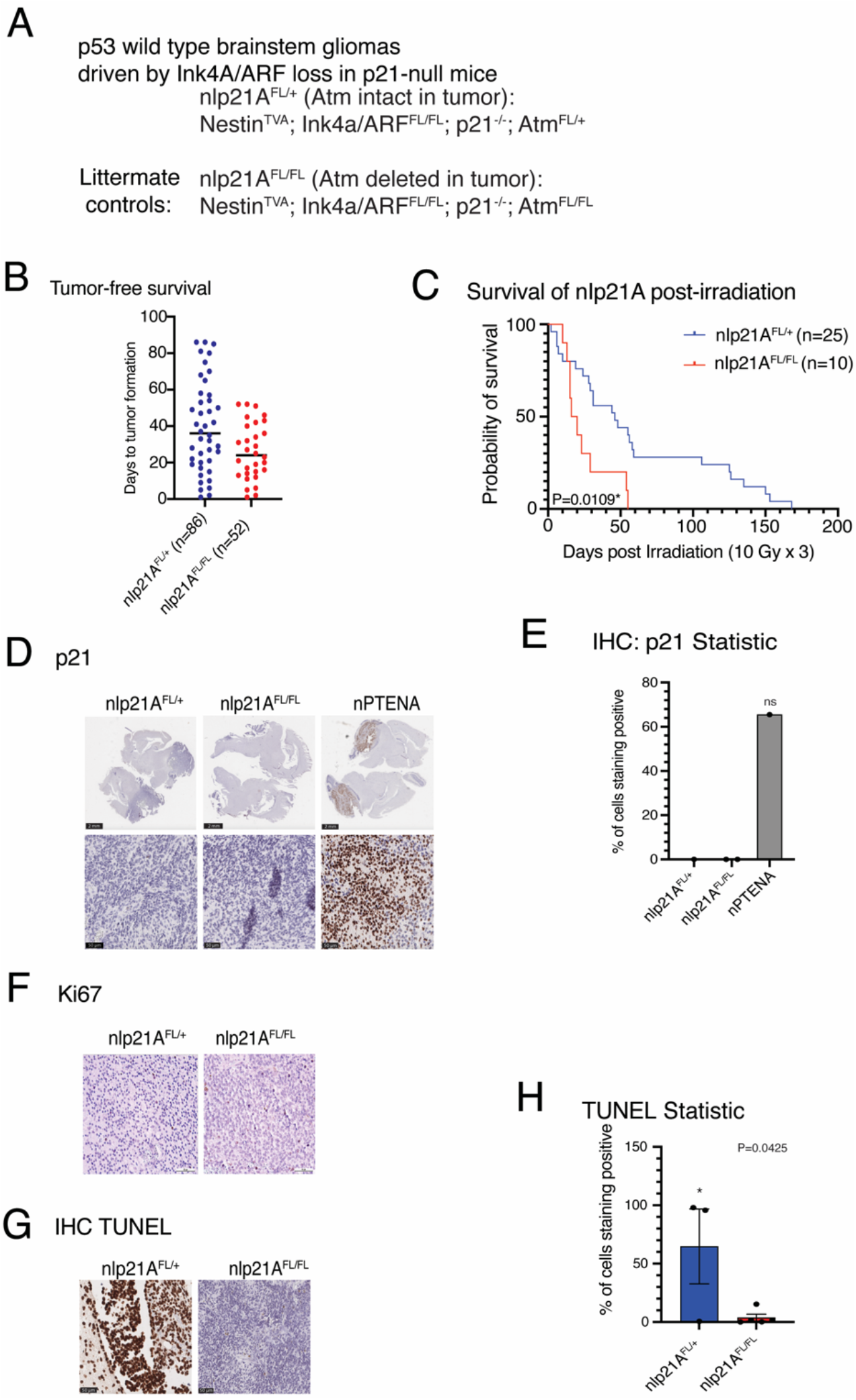
Effect of tumor-specific *Atm* loss in primary DMGs in a *Cdkn1a*-null background (p21-/-) (A) Overview of p21-/- genotypes analyzed (B) Tumor free survival of nIp21A mice with and without intact *Atm* using log-rank test. (C) Post-focal brain irradiation survival of nIp21A mice with and without intact *Atm* indicating a statistically significant survival benefit in nlp21A ^FL/+^ mice (P-value < 0.05) using log-rank test. (D) IHC showing p21 expression in nIp21A mouse brains. Nestin^TVA^; Pten^FL/FL^; Atm^FL/+^ (nPtenA) tumor-bearing brain generated with identical RCAS viruses shown as control. (E) Plot indicating percentage of tumor cells staining positive for p21 compared to total cell count. (F) IHC with Ki67 showing proliferation for nlp21A^FL/+^ and nIp21A^FL/FL^ (G) TUNEL staining of tumor-bearing brains of nIp21A mice with and without intact *Atm* in tumors collected one hour post-focal brain irradiation. (H) Quantification of TUNEL staining in nlp21A ^FL/+^ mice (P-value < 0.05) based on unpaired t-test. Scale bar = 50uM on IHC.

### A p53 transactivation domain mutant retains tumor suppressor function in mouse DMG

Since *Atm* loss could not radiosensitize *Cdkn1a*-null DMGs, we reasoned that regulation of p53 transcriptional targets other than *Cdkn1a* may cause radioresistance in p53 wild type, *Atm*-null DMGs. To investigate this possibility, we leveraged a conditional *loxP-Stop-loxP-p53*^25, 26^ allele (p53^LSL-25,26^)^33^. In the presence of Cre recombinase, this allele expresses p53^25,26^, a p53 mutant that is severely compromised for transactivation of most p53 target genes and cannot induce G1-arrest or apoptosis in response to acute DNA damage^33^. Interestingly, p53^25, 26^ retains tumor suppressor activity in lung tumors^33^, but it is unknown whether it retains tumor suppressor activity for brain tumors. We first determined if p53^25, 26^ retained tumor suppressor activity in DMG. To test this, we compared littermate mice with either p53^LSL-25,26/FL^ or p53^FL/FL^. All mice harbored Nestin^TVA^ and were injected with Cre, luciferase, and PDGF-B retrovirus constructs as described above (Figure 7A). We noted significant delay to tumor presentation in the p53 ^LSL-25,26/FL^ group compared to the p53^FL/FL^ controls (Figure 7B,C). Immunohistochemical analysis revealed heterogeneous p53 expression in the p53^LSL-25,26/FL^ group and apparently absent p53 expression in the p53^FL/FL^ group (Figure 7D,E). Thus, a p53 mutant with severely compromised transactivation activity retains tumor suppressor activity in primary mouse brainstem gliomas. These results indicate that p53 transactivation functions are dispensable for p53 tumor suppression in DMG.

**Figure 7.**
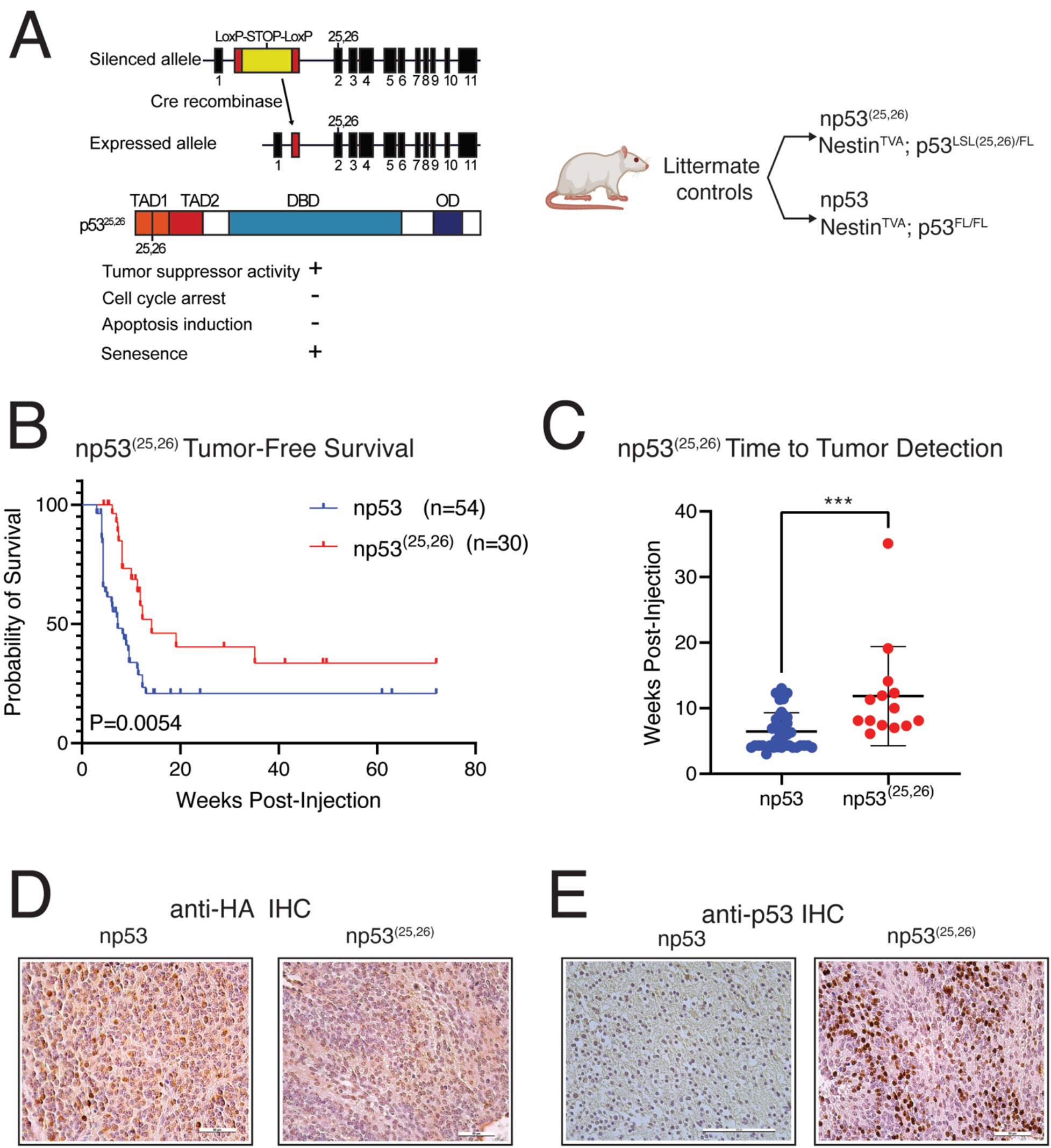
Tumor formation in mice expressing a p53 transactivation domain 1 mutant. (A) Schematic for conditional p53 transactivation domain 1 mutant, and mice genotypes for expression of a p53 transactivation domain 1 mutant. (B) (C) Tumor free survival in the np53^(25, 26)^ compared to the np53 control based on log-rank test. Time to tumor presentation in the p53^25,26/FL^ group compared to the p53-FL/FL controls with Wilcoxon test. (D) IHC for Anti-HA in p53 and p53^(25,26)^ group. Scale bar = 50 uM (E) IHC for p53 expression in p53 (Scale bar = 100 uM) and p53^(25,26)^ (50 uM) group.

### *Atm* loss does not radiosensitize mouse DMGs lacking a functional p53 transactivation domain

We next sought to determine if *Atm* loss could radiosensitize DMGs lacking p53 transcriptional activity but retaining other non-transcriptional functions of p53. We previously showed that *Atm* loss did not radiosensitize brainstem gliomas driven by Ink4A/ARF loss, but that *Atm* loss did modestly radiosensitize brainstem gliomas with both Ink4A/ARF loss and p53 loss ^5^. We reasoned that if loss of p53 transactivation domain function is the determinant of radiosensitization by *Atm* loss, then brainstem gliomas with both Ink4A/ARF loss and expression of a transactivation-deficient p53^25,26^ allele would be radiosensitized by *Atm* loss. To test if mouse DMGs with p53^25, 26^ and Ink4A/ARF loss are radiosensitized by *Atm* loss, we bred mice of genotype Nestin^TVA^; p53^LSL-25,26/FL^; Ink4A/ARF^FL/FL^; Atm^FL/FL^. To test the effects of *Atm* loss, we compared these to littermate controls with the same genotype except an intact *Atm* allele (Nestin^TVA^; p53^LSL-25,26/FL^; Ink4A/ARF^FL/FL^; Atm^FL/+^) (Figure 8A). We noted similar time to tumor formation in both models (Figure 8D. *Atm* loss was associated with differential staining of phospho-ATM and phospho-KAP1 after focal brain irradiation (Figure 8B and 8C), confirming loss of ATM functional activity. After subjecting the mice to fractionated focal brain irradiation, no difference in overall survival was appreciated (Figure 8E. These results indicate that transactivation-independent functions of p53 may be the primary determinants of whether mouse DMGs can be radiosensitized by *Atm* loss.

**Figure 8.**
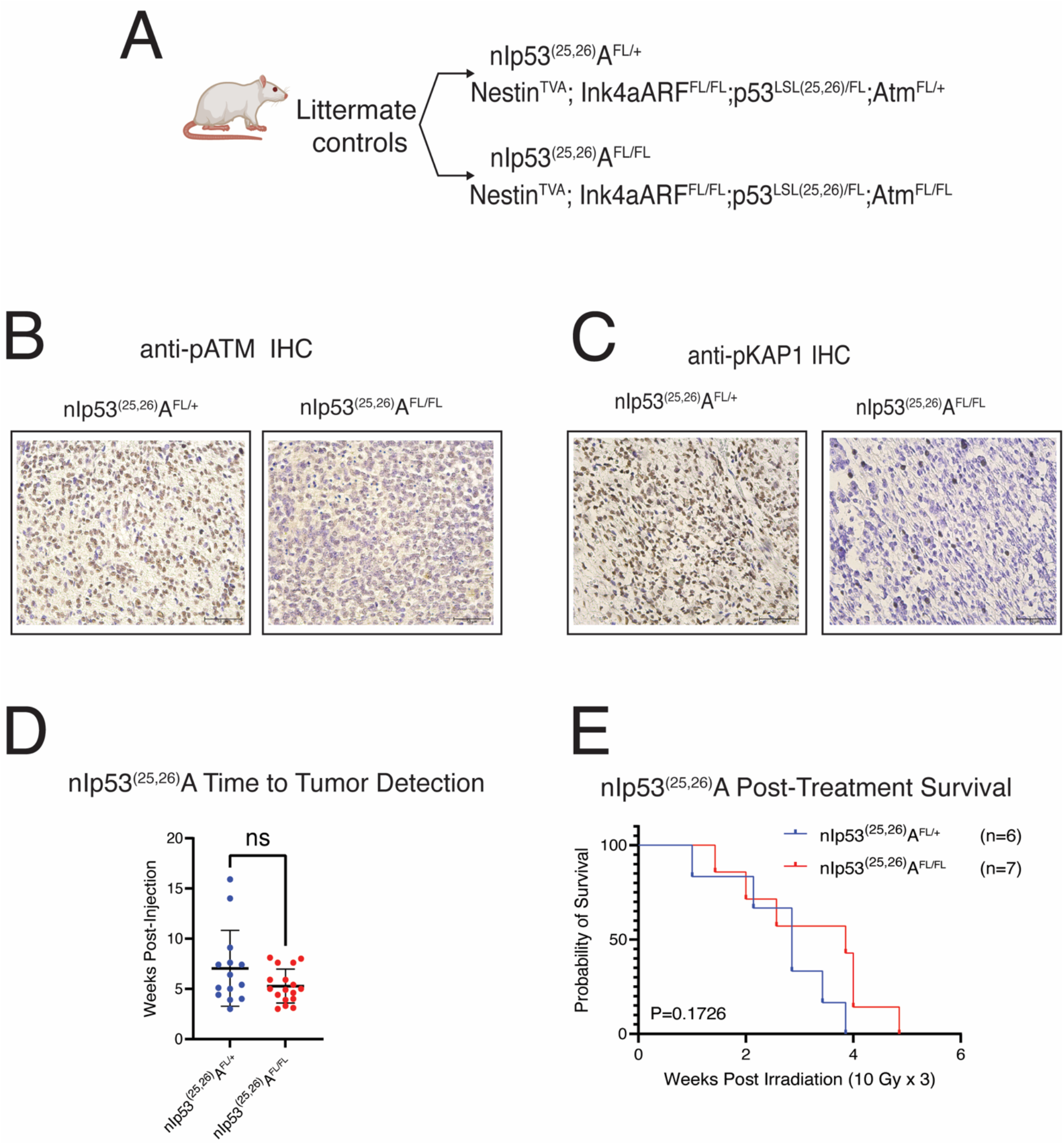
Effect of *Atm* loss on survival after fractionated focal brain irradiation in mouse DMGs expressing a p53 transactivation domain 1 mutant. (A) Schematic showing p53^LSL(25,26)^ allele and genotypes with Nestin^TVA^; p53^LSL(25,26)/FL^; Ink4A/ARF^FL/FL^ mice with either Atm^FL/FL^ or Atm^FL/+^ (B) IHC showing phosphor-Atm in Atm ^FL/+^ and Atm^FL/FL^ tumors. Scale bar = 50uM. (C) IHC showing phosphor-Kap1 expression in Atm^FL/+^ and Atm^FL/FL^ tumors. Scale bar = 50 uM. (D) Time to tumor formation in Nestin^TVA^; p53^LSL(25,26)/FL^; Ink4A/ARF^FL/FL^ mice with either Atm^FL/FL^ or Atm^FL/+^ (dot plot). ns, no statistical significance by Wilcoxon test. (E) Overall survival following fractionated brain irradiation in mouse DMGs expressing a p53 transactivation domain 1 mutation with or with *Atm* loss, with P-value based on log rank test.

## Discussion

Here, we describe the generation of primary mouse DMGs leveraging recent advances in murine genetic engineering including a conditional H3.3K27M allele and the RCAS/tv-a retrovirus platform ^10, 13, 14^. We use the model to show that genetic loss of *Atm*, an important target for drugs that have entered clinical trials for brain tumor patients^18^, radiosensitizes primary DMG models. Our results in p53-null/H3.3K27M mouse DMGs are similar to those reported previously for p53-null mouse brainstem gliomas^5^, in which the sole difference is the presence of H3.3K27M expression from the endogenous *H3f3a* locus in the neoplastic tumor cells in our current model. We have generated several unique genetically engineered mouse models with differential responses based on genotype (Table 1). The results from our models suggest that H3.3K27M is not a primary determinant of tumor radiosensitivity, or on the ability for targeting ATM to enhance the efficacy of radiation therapy in primary mouse DMG models.

**Table 1.** Summary of the effect of Atm loss on radiation sensitivity in genetically engineered DMG mouse models.

| Genotype<br>(Nestin<sup>TVA</sup> + Cre, Luc, PDGFB) | Baseline Radiation<br>Therapy sensitivity | Effect of *Atm* Loss on response to<br>radiation therapy |
| --- | --- | --- |
| p53<sup>FL/FL</sup> | Resistant | More sensitive<sup>5</sup> |
| Ink4A/ARF<sup>FL/FL</sup> | Sensitive | No effect<sup>5</sup> |
| Pten<sup>FL/FL</sup> | Sensitive | No effect<sup>11</sup> |
| p53<sup>FL/FL</sup>; Ink4A/ARF<sup>FL/FL</sup> | Very Resistant | More sensitive<sup>5</sup> |
| H3f3a<sup>LSL-K27m/+</sup>; p53<sup>FL/FL</sup> | Resistant | More sensitive |
| p21<sup>-/-</sup>; Ink4A/ARF<sup>FL/FL</sup> | Sensitive | More resistant |
| p53<sup>LSL-25,26/FL</sup>; Ink4A/ARF<sup>FL/FL</sup> | Resistant | No effect |

Our results from genetic experiments in primary mouse models nominate p53 as a key determinant for the ability for DMG to be radiosensitized by *Atm* loss. Almost all p53-altered primary mouse models are radioresistant and are radiosensitized by *Atm* loss, including (i) a model driven by p53 loss with wild type *H3f3a*^5^, (ii) a model driven by both p53 loss and by loss of Ink4A/ARF^5^, and (iii) the H3.3K27M/TP53 mutant model reported here (Figure 2C). In contrast, *Atm* loss is unable to radiosensitize primary p53-wildtype brainstem glioma mouse models, including models driven by Ink4A/ARF loss^5^ and models driven by *Pten* loss^11^. Of note, a recent study comprehensively profiled that pharmacologic ATM inhibition radiosensitized both p53-mutant and p53-wild type patient-derived models of DMG and pediatric high-grade glioma^6^. Together, these data suggest that p53 mutational status should be tested in correlative analyses for future clinical trials of ATM inhibitors in DMG patients.

Our ISS data provides the first high-resolution transcriptional analysis at high gene plexy (∼300 gene targets) of a mouse tumor model. High-resolution spatially resolved transcriptomic profiling was especially critical in this model since vasculature and neoplastic compartments within the tumor play distinct roles in therapeutic response^12^. Future work will leverage these data to interrogate tumor immune and vascular microenvironment alterations induced by irradiation and *Atm* loss, which may guide the rational design of combinations between radiation therapy, ATM inhibitors, and therapies targeted at the immune system or vasculature.

The current work implicates transactivation-independent mechanisms by which p53 mediates radioresistance in *Atm*-null tumors. Our ISS data showed that irradiation elicited overexpression of *Cdkn1a*, a key downstream target of p53 that mediates cell senescence and G1-to-S checkpoint arrest, in p53-null tumors suggesting p53-independent mechanisms of *Cdkn1a* expression^34^. This finding led us to dissect the contribution of p53 transactivation functions that regulate p21 expression to radiosensitization in *Atm*-null DMGs. While *Atm* loss radiosensitizes tumors lacking p53, we found that *Atm* loss could not radiosensitize tumors containing a p53^25, 26^ allele deficient in p53 transactivation function. Similarly, tumors that lacked *Cdkn1a* (p21) could not be radiosensitized by *Atm* loss. Our findings highlight the importance of carefully considering the p21 status in clinical trials involving Atm inhibition given the complex role of p21 in tumor growth and microenvironment^35^. Strikingly, we found that *Atm* loss made the tumors more radioresistant in mice that lacked *Cdkn1a*, and that this finding was associated with increased apoptosis. These findings implicate transactivation-independent functions of p53 as key determinants of radiosensitivity in *Atm*-null tumors. Future work will dissect transactivation-independent functions of p53 such as promoting apoptosis through mitochondrial membrane permeabilization, directly repressing transcription, and/or directly interacting with complexes that detect DNA lesions^36, 37^. Our data provides genetic and mechanistic insight that builds upon studies of pharmacologic ATM inhibition in patient derived xenograft models^6^. Further studies are needed to determine transactivation-independent mechanisms of p53 and ATM-directed therapies and their impact on overcoming resistance to radiation therapy in patients with H3.3K27M-mutant DMG.

## Data and Code Availability

In situ sequencing data have been deposited at GEO and will be publicly available as of the date of publication. Accession numbers will be listed in the key resources table. This paper does not report original code. Any additional information required to reanalyze the data reported in this paper is available from the lead contact upon request.

## Acknowledgements

We thank sources of funding including NCI K08256045 Mentored Clinician Scientist Development Award, ChadTough Defeat DIPG Foundation, the SoSo Strong Foundation, the Pediatric Brain Tumor Foundation, the Emily Beazley’s Kures for Kids Fund, the St. Baldrick’s Foundation, Lauren Brescia Memorial Fund, and NCI P50CA190991 Duke SPORE in Brain Cancer developmental funds to ZJR. This work was also supported by 7R35CA197616 from the NCI to DGK. We thank Dr. Suzanne Baker for the gift of the H3f3a-loxP-Stop-loxP-K27M-Tag mice. Funds for ISS data generation were provided by the Duke Brain Tumor Omics Program.

## Author contributions

AM, ZJR, SGG, DGK, DMA and OJB devised the study and designed experiments. SW, HL, BEF, LW, MEG, DG, LL, KD, and ZJR performed mouse experiments and tabulated data. NTW performed mouse irradiations. KA and NH performed patient-derived xenograft and ATM inhibitor pharmacologic experiments. LA contributed p53 transactivation mutant mouse strain and experimental design regarding the strain. EH, KA, LW, and ZJR assisted with in situ sequencing experiments. MA, VJ, JAR, and ZJR performed bioinformatic analyses. AM, ZJR, and SGG prepared the manuscript.

## Inclusion and diversity

One or more of the authors of this paper self-identifies as an underrepresented ethnic minority in their field of research or within their geographical location. One or more of the authors of this paper self-identifies as a gender minority in their field of research.

## Methods

All animal experiments performed were approved by the IACUC at the individual institutions. Detailed workflows for generation, brain irradiation, and molecular analysis of primary mouse DMG models using RCAS/tv-a and Cre/loxP technologies are found in our recent *STAR Protocols* manuscript^38^. All new reagents, materials, and software are listed in the key resources table below.

### Mouse strains

Detailed workflows for generation, brain irradiation, and molecular analysis of primary mouse DMG models using RCAS/tv-a and Cre/loxP technologies are found in our recent *STAR Protocols* manuscript^38^. Complex mouse strains were generated by breeding mice with the following alleles together: Nes^TVA^, Atm^FL^, and p53^FL^ alleles^38^, Ink4A/ARF^FL^ allele^38^, the p53^LSL(25,26)^ allele^33^, and the p21^-/-^ allele^39^. The H3f3a^LSL-K27M-Tag^ allele is a gift from Dr. Suzanne Baker^10^.

### DF1 cell culture and retrovirus generation

DF1 cells were cultured in DMEM media containing 10% FBS and RCAS/tv-a retroviruses were generated utilizing RCAS-Cre, RCAS-luc, and RCAS-PDGFB plasmids as previously described^38^.

### Mouse brainstem injection

The harvested DF1 cells were injected into brainstems of mice anesthetized on ice at postnatal day 4 as described^38^. All institutional approvals were obtained prior to injection. Patient-derived xenografts using the SF8628 model were generated by brainstem injection as described^20–22^.

### Mouse *in-vivo* imaging

The bioluminescence imaging of gliomas within mice was performed utilizing intraperitoneal injection of D-luciferin and an IVIS Illumina III as previously described^38^.

### Image Guided focal brain irradiation

Irradiation to gliomas within mice was delivered on a Small Animal Radiation Research Platform (SARRP) using image-guided, opposed-lateral beams as described^38^. 10 Gy times three consecutive daily fractions was delivered for primary DMG models. 2 Gy times three days a week to 12 Gy total was delivered for patient-derived xenograft DMG models.

### ATM inhibitor studies

SF8628 (H3.3K27M DIPG) was obtained from the University of California San Francisco (UCSF) medical center, and in accord with an institutionally approved protocol. Establishment of SF8628 cell culture from surgical specimens, and tumor cell modification for expression of firefly luciferase for in vivo bioluminescence imaging, have been described^20–22^. The SF8628 cells were propagated as monolayers in complete medium consisting of Dulbecco’s Modified Eagle’s medium (DMEM) supplemented with 10% fetal bovine serum (FBS, A3160) and non-essential amino acids. Short tandem repeats (STR) were obtained to confirm the identity of the cell lines. All cells were cultured in an incubator at 37°C in a humidified atmosphere containing 95% O2 and 5% CO2 and were mycoplasma-free at the time of testing. Six-week-old female athymic mice (rnu/rnu genotype, BALB/c background) were purchased from Envigo and housed under aseptic conditions. Pontine injection of tumor cells was performed as previously described^20–22^. Each mouse was injected with 1 μL of SF8628 cell suspension (100,000 cells/μL) into the pontine tegmentum at a depth of 5 mm from the inner base of the skull. For the efficacy study of AZD1390 and radiation, animals were randomized into four treatment groups: 1) vehicle control (0.5% hydroxymethylcellulose, 0.1% Tween 80, n=6), 2) ADZ1390 monotherapy (oral gavage of 20 mg/kg of AZD1390 for 5 times a week for two consecutive weeks, n=6), 3) radiation monotherapy (2.0 Gy, 3 times a week for two consecutive weeks for a total dose of 12 Gy, n=6), 4) AZD1390 and radiation combination therapy (n=6). Biweekly bioluminescence imaging was used to monitor tumor growth and response to therapy as previously described^20^. Mice were monitored daily and euthanized at endpoint which included irreversible neurological deficit or body condition score less than 2. All animal protocols were approved by the Northwestern University Institutional Animal Care and Use Committee. AZD1390 was obtained from AstraZeneca.

### Immunohistochemistry

IHC was performed utilizing previously described methods for Ki67, p-Atm, p-KAP1, and p53^38^. Additional IHC and TUNEL staining was performed by HistoWiz Inc. (histowiz.com) using a Standard Operating Procedure and fully automated workflow for p21 (Cdkn1a), FLAG, and TUNEL on a Bond Rx autostainer (Leica Biosystems) with enzyme treatment (1:1000) using standard protocols.

### Quantification and Statistical Analysis

Statistical significance in Volcano plots across ROIs was assessed with the Wilcoxon test on SCTransform normalized count data using the FindMarkers Seurat function. Data plotting and quantification analysis were performed using GraphPad Prism v.9. Unpaired t-test was utilized to determine significance in quantification on IHC. Log-Rank test was utilized to determine survival. Wilcoxon test was used to determine differences in time to tumor detection. Individual data points were plotted, and all statistically significant values (p value less than 0.05) are identified with an asterisk (*).

### Xenium In Situ and Bioinformatics Analysis

Tumor-bearing brains subjected to Xenium ISS were detected by *in vivo* imaging 37-48 days after birth and collected seven days after tumor detection, either after three daily treatments of 10 Gy initiated within two days of tumor detection, or mock treatment. Initial data generated by Xenium instrument^40^ is processed on board with a built-in analysis tool called Xenium Analyzer^40^. The Xenium Analyzer is fully automated and includes an imager (imageable area of about 12 x 24 mm per slide), sample handling, liquid handling, wide-field epifluorescence imaging, capacity for two slides per run, and an on-instrument analysis pipeline. The analysis pipeline includes image pre-processing, puncta detection, transcript decoding and quality score assignment. The pipeline also performs cell segmentation using DAPI images to detect nuclei using a neural network. Then each nucleus is expanded outwards until either 15 um max distance is reached or the boundary of another cell is reached. A variety of output files are produced by the on-instrument pipeline. The essential files for downstream analysis include the feature-cell matrix (HDF5 and MEX formats identical to those output by single cell RNA tools from 10X (Cellranger/Spaceranger), the transcripts (listing each mRNA, its 3D coordinates, and a quality score), and the cell boundaries CSV file.

Specific regions of the tissue were annotated manually using the polygon tool in the Xenium Explorer software (development version, 10x Genomics), and the polygon coordinates were exported as csv files.

Xenium output was first imported with R (4.3.1) using the LoadXenium function from Seurat (4.9.9.9050)^41^. Cells with zero counts were then filtered, and the points within the polygon coordinates were identified using the point.in.polygon function from the sp (2.0.0) R package. Further plots were generated using Seurat, and deconvolution was performed with spacexr (2.2.1)^42^ using a custom annotated single cell reference from a previous experiment. DGE analysis across ROIs was assessed with the Wilcoxon test on SCTransform normalized count data using the FindMarkers Seurat function. Co-occurrence and neighborhood enrichment plots were made using the Python package Squidpy (1.2.3)^43^, and trajectory and cell-cell-interaction analysis was performed using the Python package STLearn (0.4.12) ^44^.

All cells on the entire slide were utilized to determine the different types of cell cluster. All unlabeled cells were removed for the Squidpy and STLearn analysis. Co-occurrence and neighborhood enrichment analysis were conducted on all cells within the entire slide. Tumor core and periphery were utilized to identify the Cell:Cell and Cell:Ligand interactions. Differentially expressed gene analyses were conducted on tumor core.

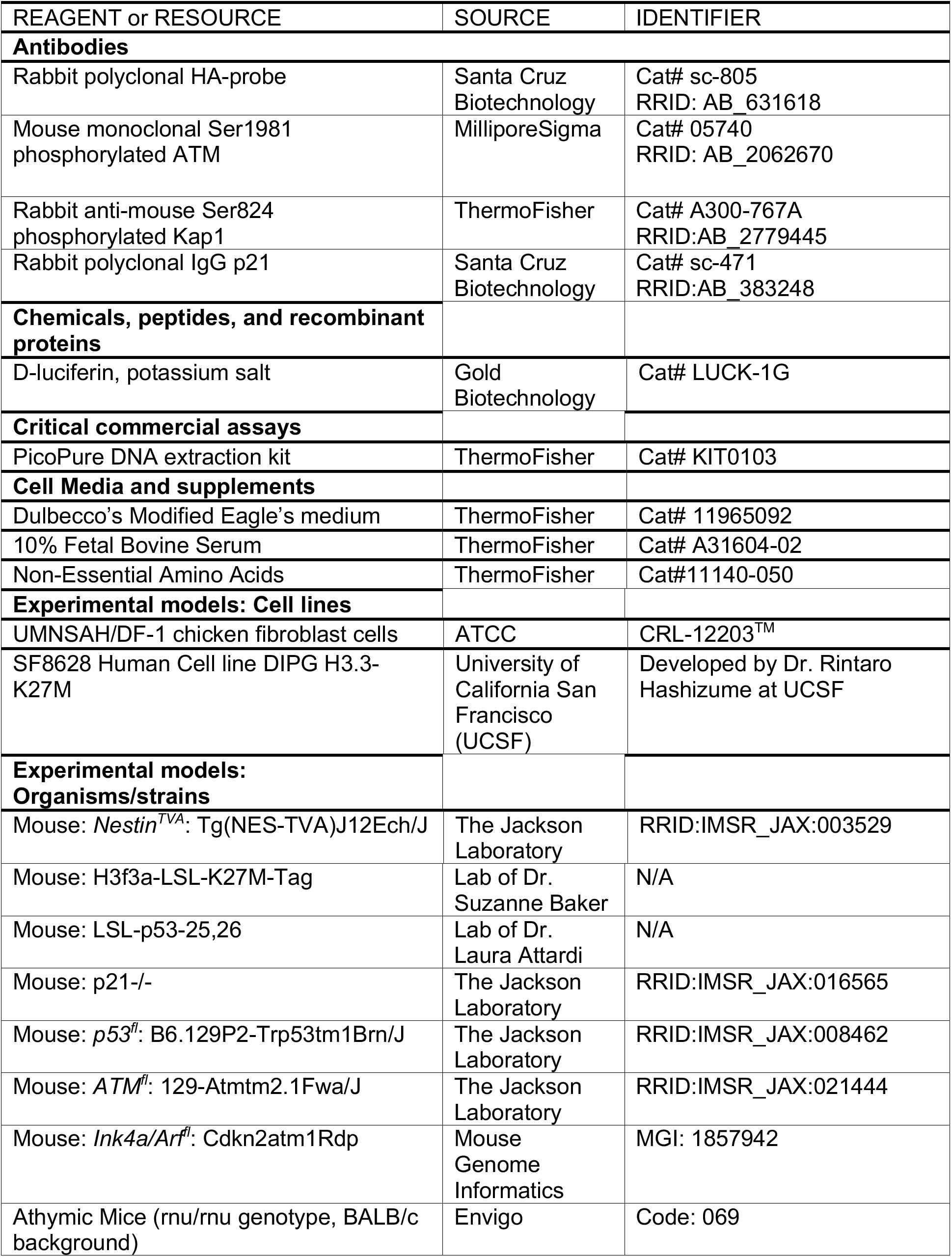
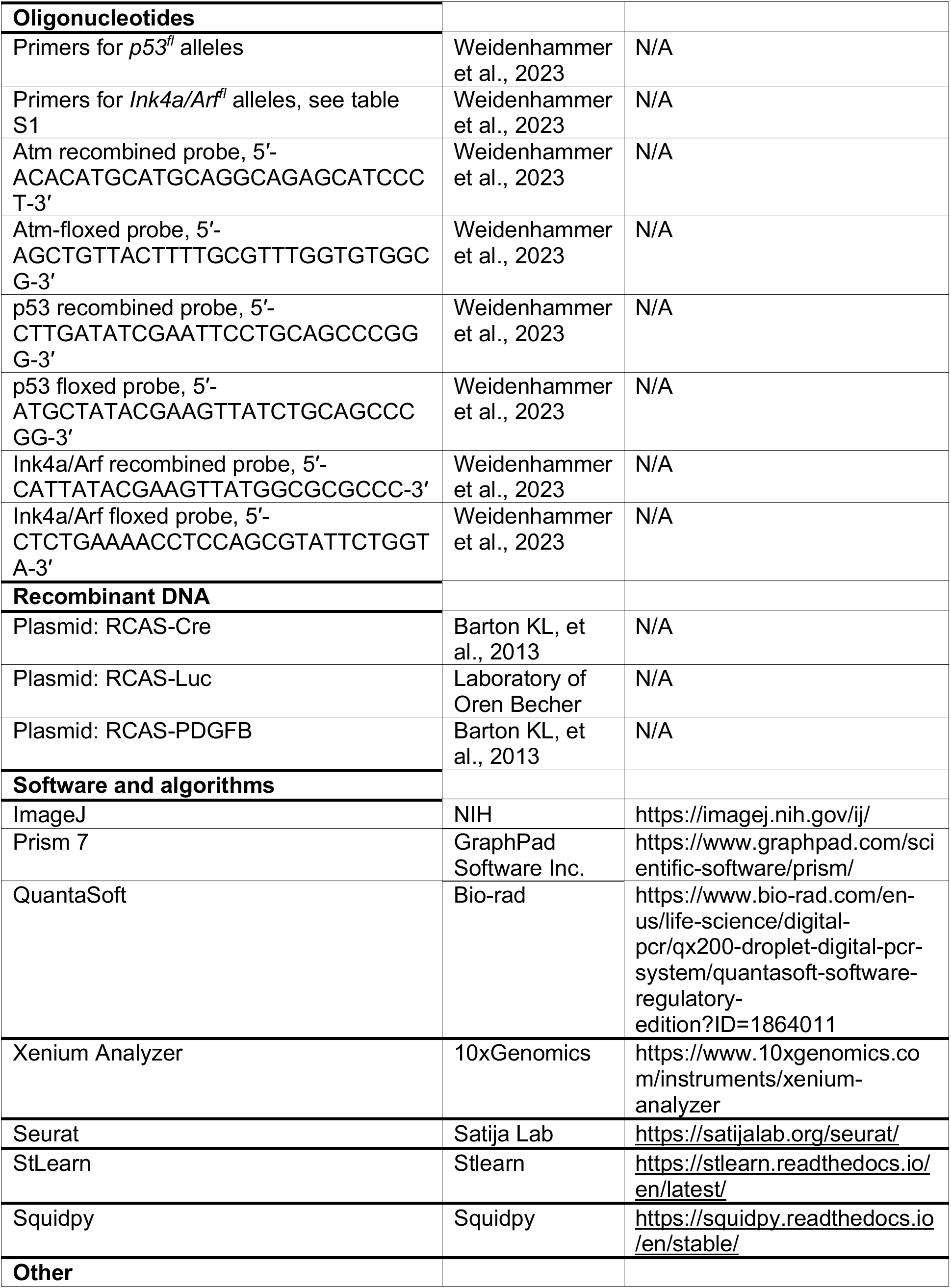
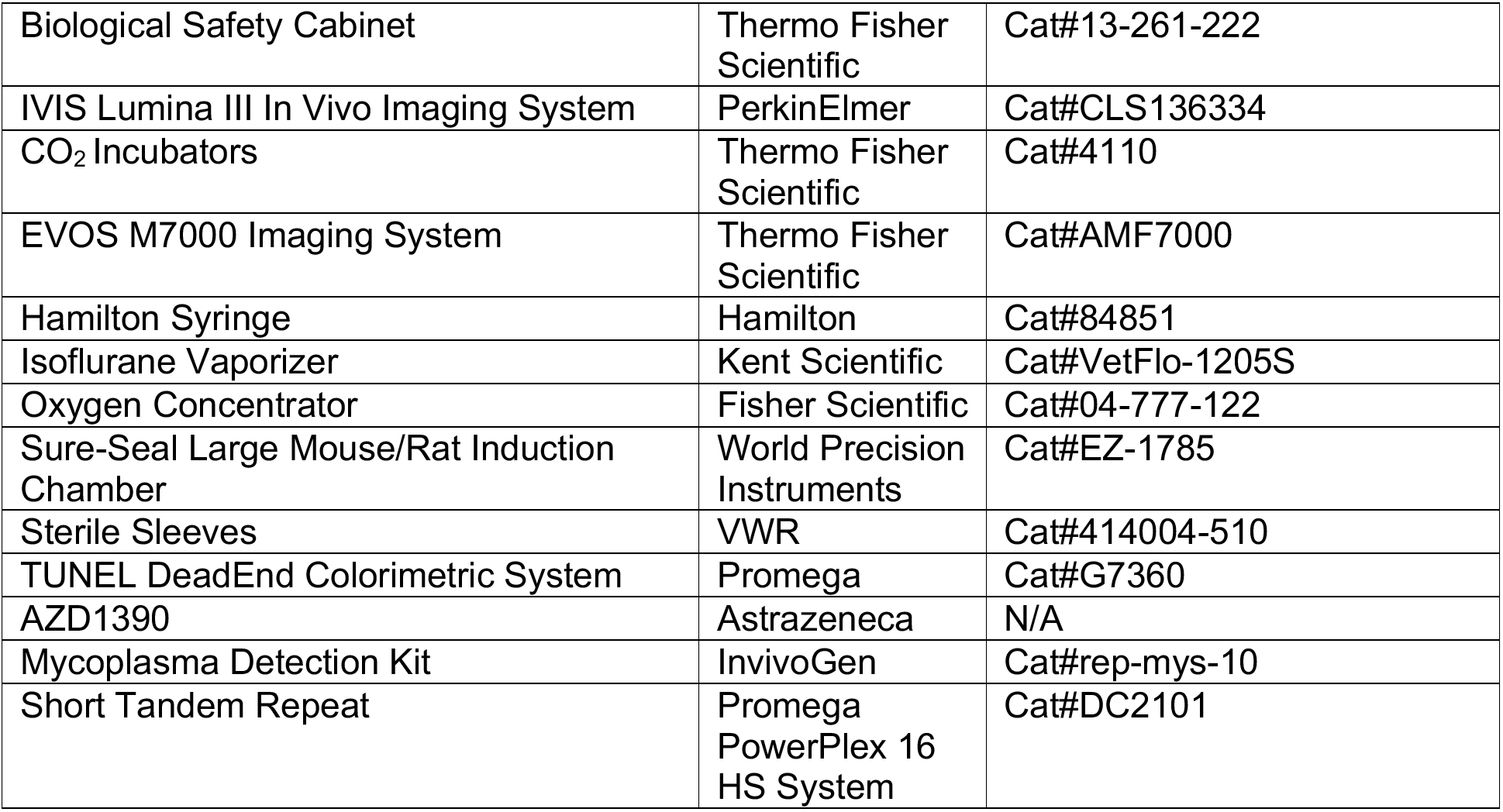
Key resources table.

### Materials and equipment

D-Luciferin Stock Solution.

Materials and equipment D-Luciferin Stock Solution
| Reagent | Final concentration | Amount |
| --- | --- | --- |
| D-Luciferin, Potassium Salt | n/a | 1 g |
| Dulbecco's Phosphate Buffered Saline without Ca<sup>2+</sup> and Mg<sup>2+</sup> | 1X | 66.6 mL |
| **Total** | **15mg/mL** | **66.6 mL** |

Store at -80°C; expires after 1 year

#### Alternatives

D-Luciferin Sodium Salt and L-Luciferin Potassium Salt can be substitute for D-Luciferin, Potassium Salt

