## Supplementary Figures and Tables for "*Ataxia-telangiectasia mutated* (*Atm*) disruption sensitizes spatially-directed H3.3K27M/TP53 diffuse midline gliomas to radiation therapy"

SUPPLEMENTAL FIGURES AND TABLES

Figure S1 – UMAP Single Cell phenotyping of a representative tumor-bearing brain. (Page 2-3).

Figure S2 – Differentially expressed marker genes for each cluster identified within all tumors (n=4). (Pages 4 – 8).

Figure S3 – Collapsed individual cell clustering into 10 archetypal cell types. (Page 9).

**Supplementary Tables**

Table S1 – Panel of 298 mouse brain and DMG transcripts targeted by in situ sequencing. (10-20)

Table S2 – Collapsed individual cell clustering into 10 archetypal cell types. (Page 21).

Table S3 –Top differentially expressed genes of *Atm* intact (FL/+) with and without irradiation. (Pages 22-23).

Table S4 –Top differentially expressed genes of *Atm* null (FL/FL) with and without irradiation. (Pages 24-25).

Table S5 – Top Cell Ligand receptors with a p-value < 0.05 for all tumors. (Pages 26-27)

Figure S1 – UMAP Single Cell phenotyping of a representative tumor-bearing brain.

UMAP Cell Cluster for representative tumor sample Atm null (nPHA $^{FL/FL}$)

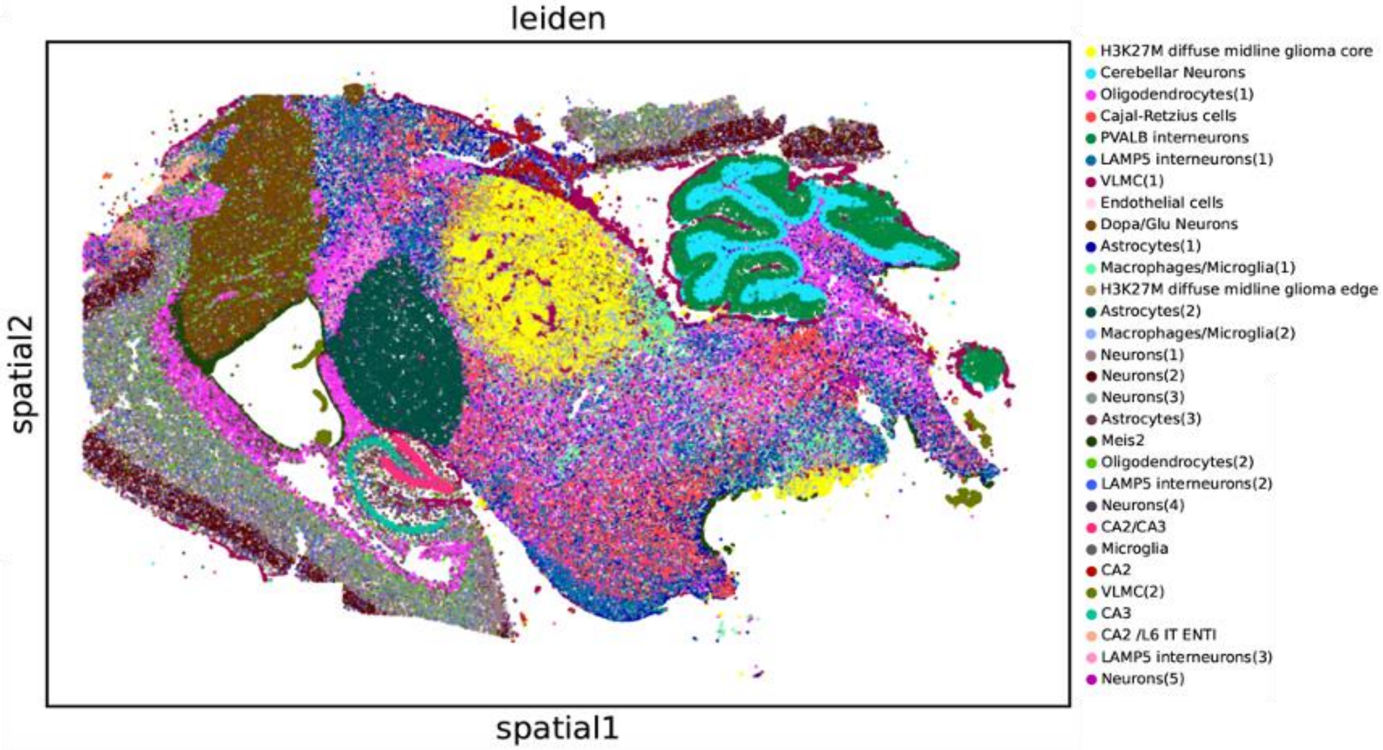


Single cell phenotyping for representative tumor sample Atm null (nPHA$^{FL/FL}$)

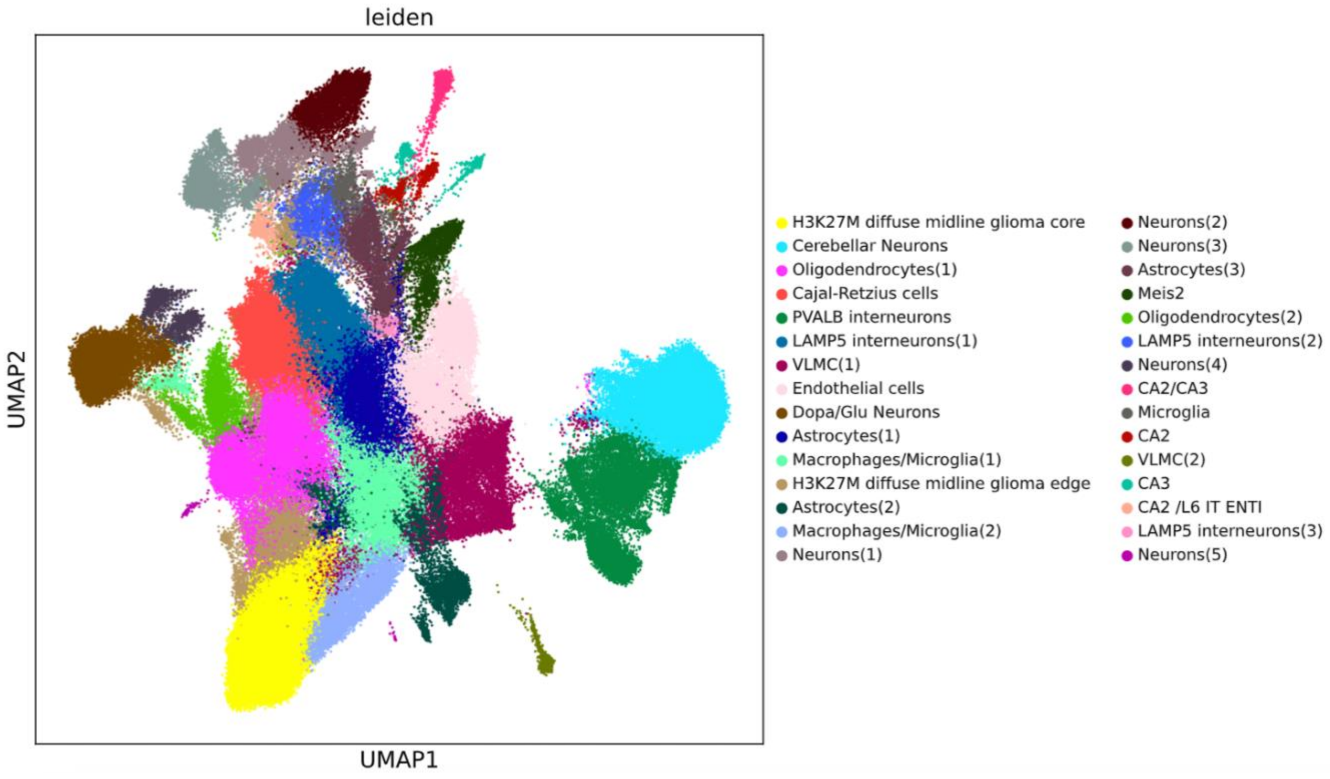

**Figure S1 - UMAP clustering of Xenium ISS from a representative tumor-bearing brain.** At top, sagittal section of an unirradiated mouse brain bearing a primary diffuse midline glioma containing H3.3K27M mutation and p53 and Atm loss (Nestin$^{TVA}$; p53$^{FL/FL}$ ; H3f3a$^{LSL\text{-}K27M\text{-}Tag}$; Atm$^{FL/FL}$; nPHA$^{FL/FL}$) shown in Figure 3B. Tumor was generated by injection of chicken fibroblast cells producing Cre, luc, and PDGFB RCAS retroviral vectors. Cell types are inferred based on differentially expressed cell markers. At bottom, UMAP clustering of cells from the same sample.

### Figure S2: Differential expressed marker genes for each cluster within each tumor

#### $Atm^{FL/+}$ ($nPHA^{FL/+}$)

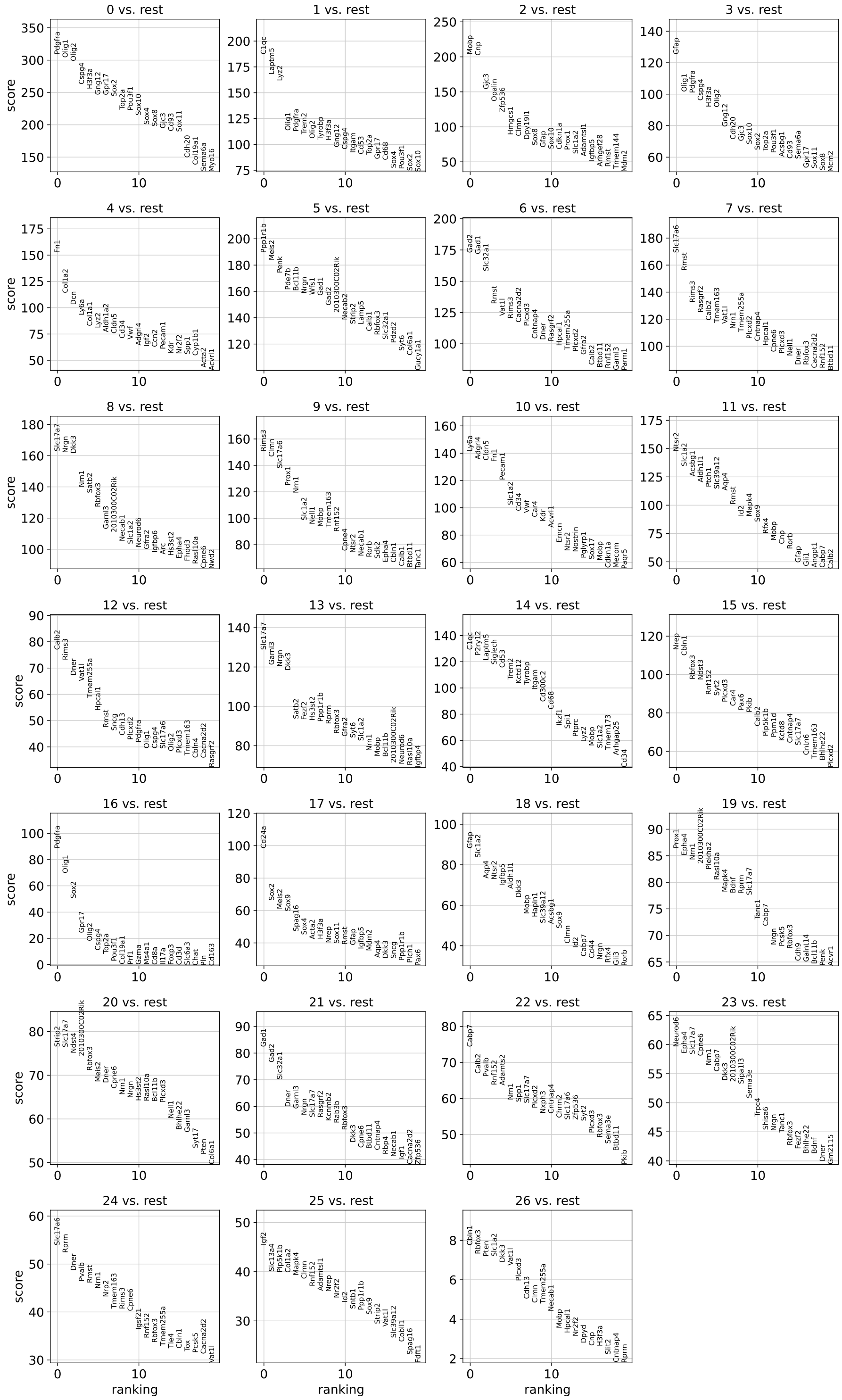

#### $Atm^{FL/+}$ ($nPHA^{FL/+}$) with irradiation (10Gy x 3)

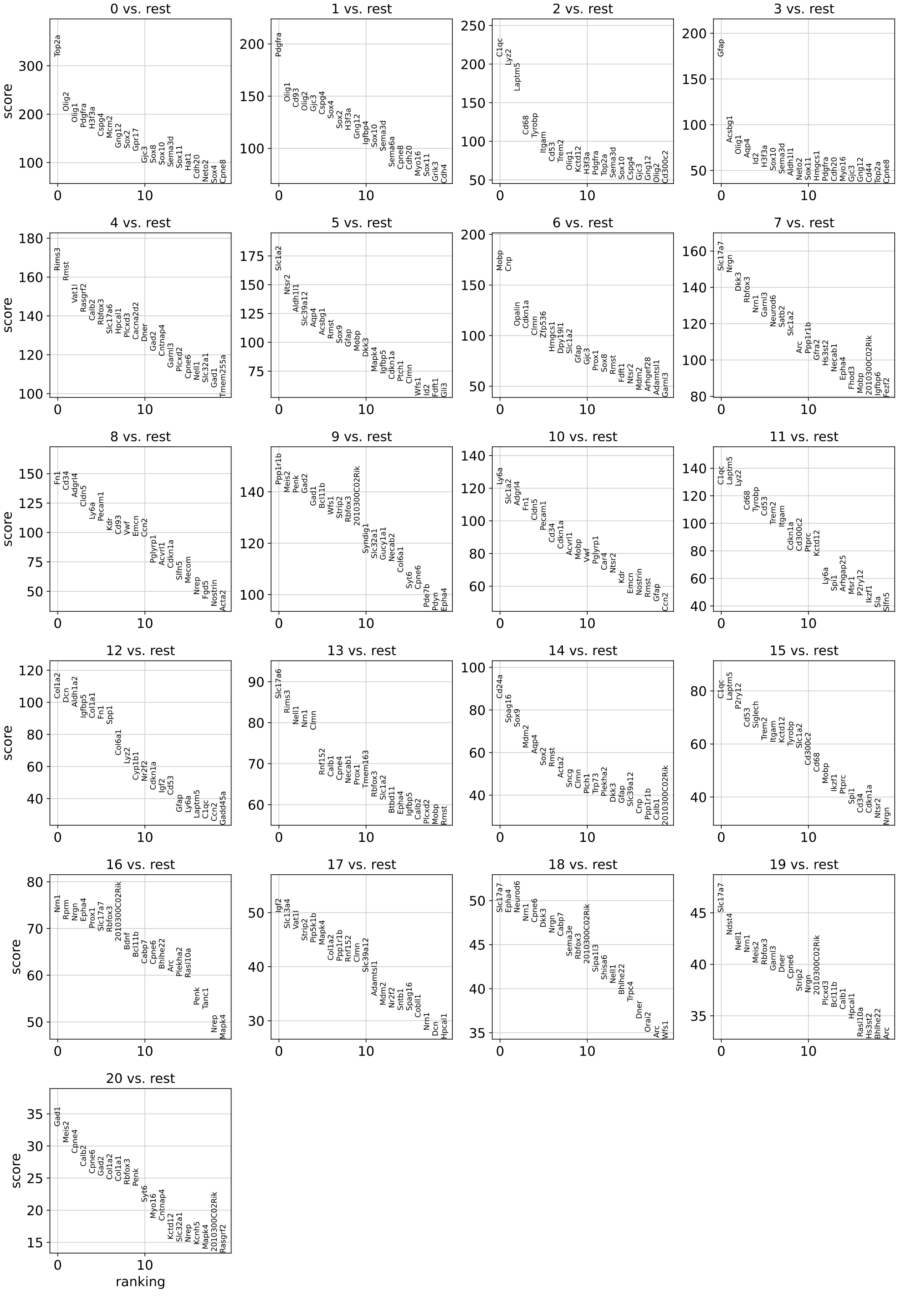

Atm$^{FL/FL}$ (nPHA$^{FL/FL}$)

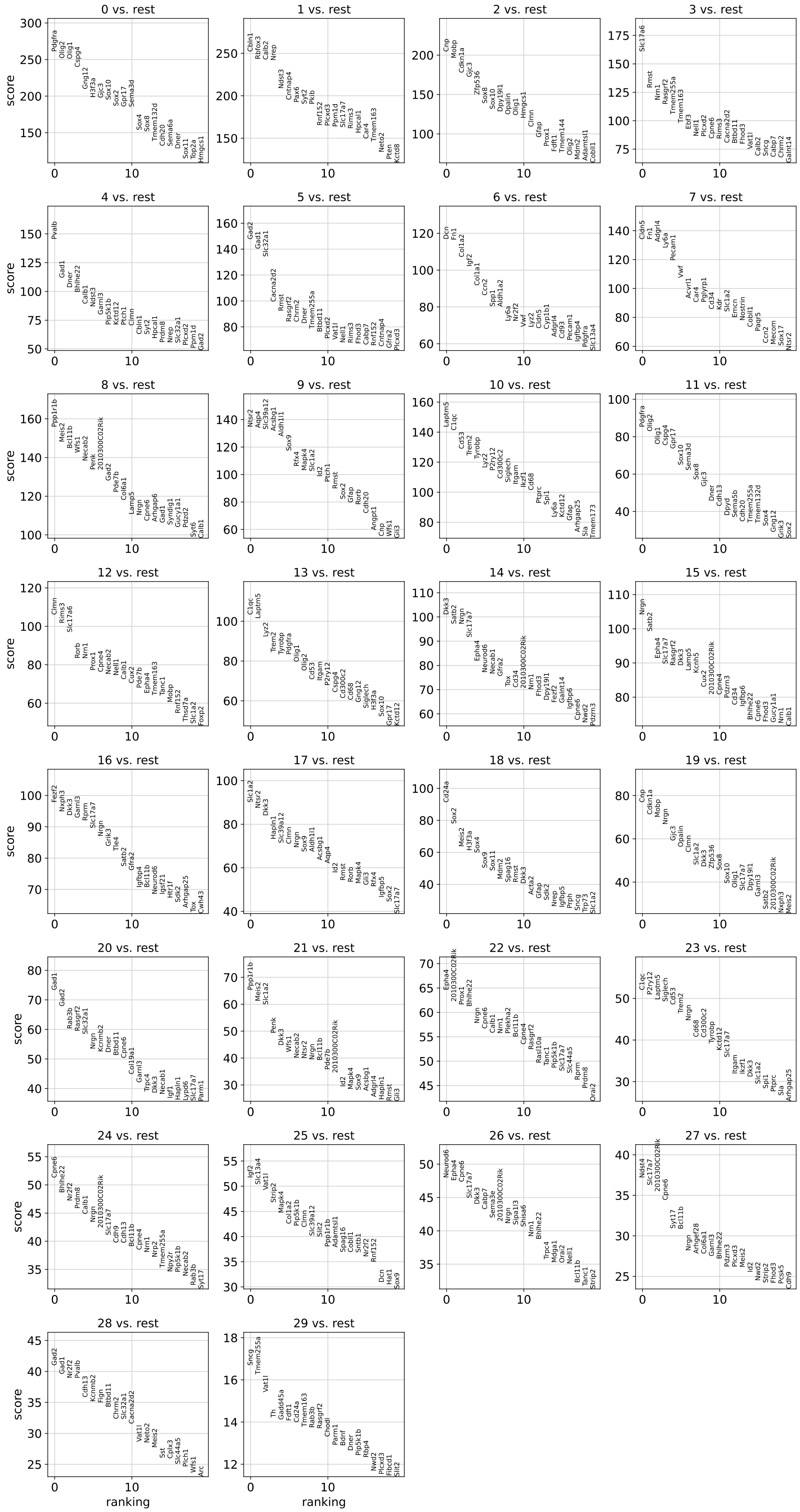

### Atm$^{FL/FL}$ (nPHA$^{FL/FL}$) with irradiation (10Gy x 3)

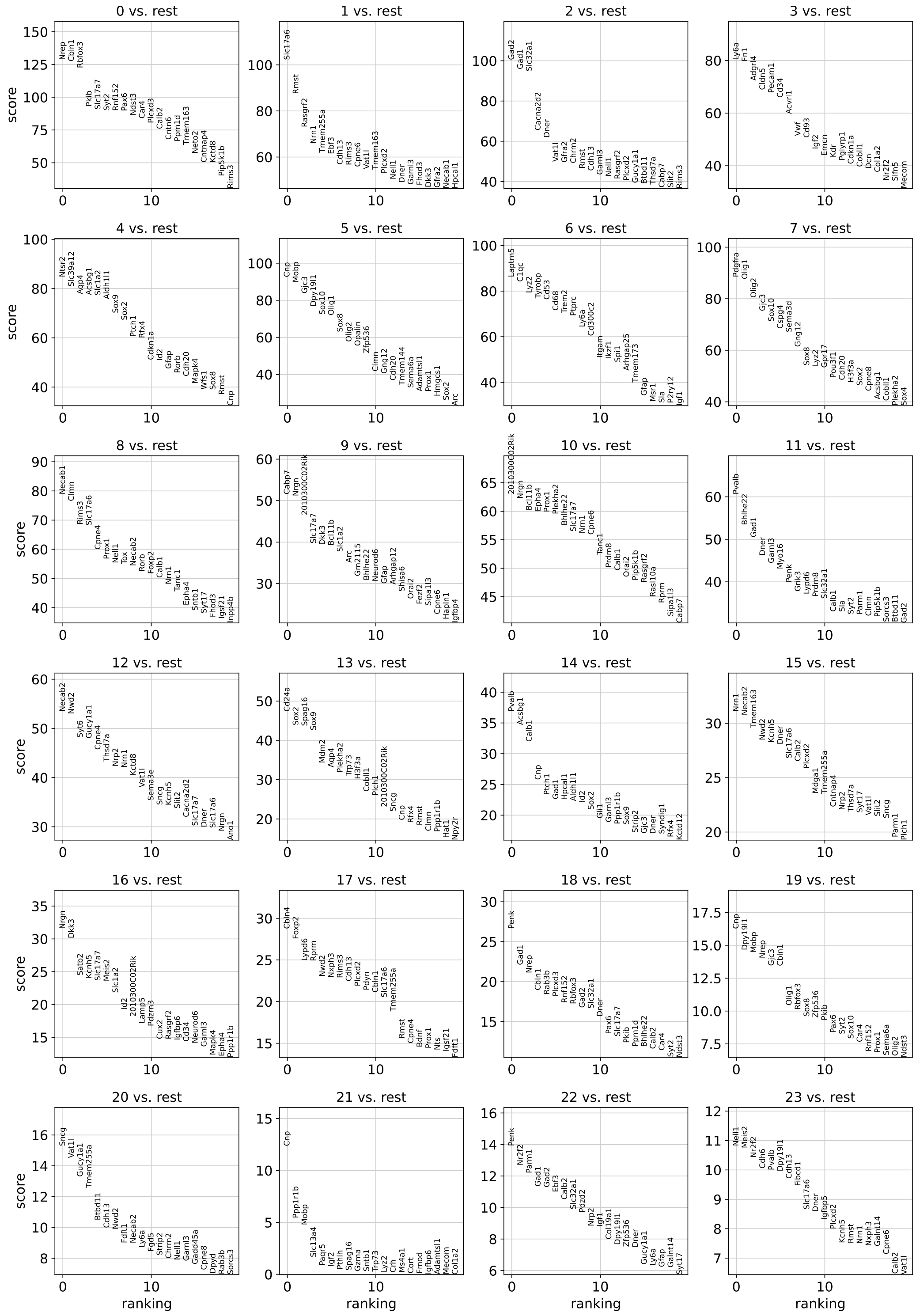

**Figure S2 - Differentially expressed marker genes for each cluster identified within all tumors.**

Plots show top differentially expressed genes for each cluster compared to all other clusters within the same sample. Four samples are shown, including $Nestin^{TVA}$; $p53^{FL/FL}$ ; $H3f3a^{LSL\text{-}K27M\text{-}Tag}$; $Atm^{FL/+}$($nPHA^{FL/+}$) without irradiation; $Nestin^{TVA}$; $p53^{FL/FL}$ ; $H3f3a^{LSL\text{-}K27M\text{-}Tag}$; $Atm^{FL/+}$ ($nPHA^{FL/+}$) status post 10 Gy x 3 focal brain irradiation; $Nestin^{TVA}$; $p53^{FL/FL}$ ; $H3f3a^{LSL\text{-}K27M\text{-}Tag}$; $Atm^{FL/+}$ ($nPHA^{FL/FL}$) without irradiation; and $Nestin^{TVA}$; $p53^{FL/FL}$ ; $H3f3a^{LSL\text{-}K27M\text{-}Tag}$; $Atm^{FL/+}$ ($nPHA^{FL/FL}$) status post 10 Gy x 3 focal brain irradiation. Score represents relative fold change and significance of increased expression compared to all other groups.

Figure S3: Cell Clustering

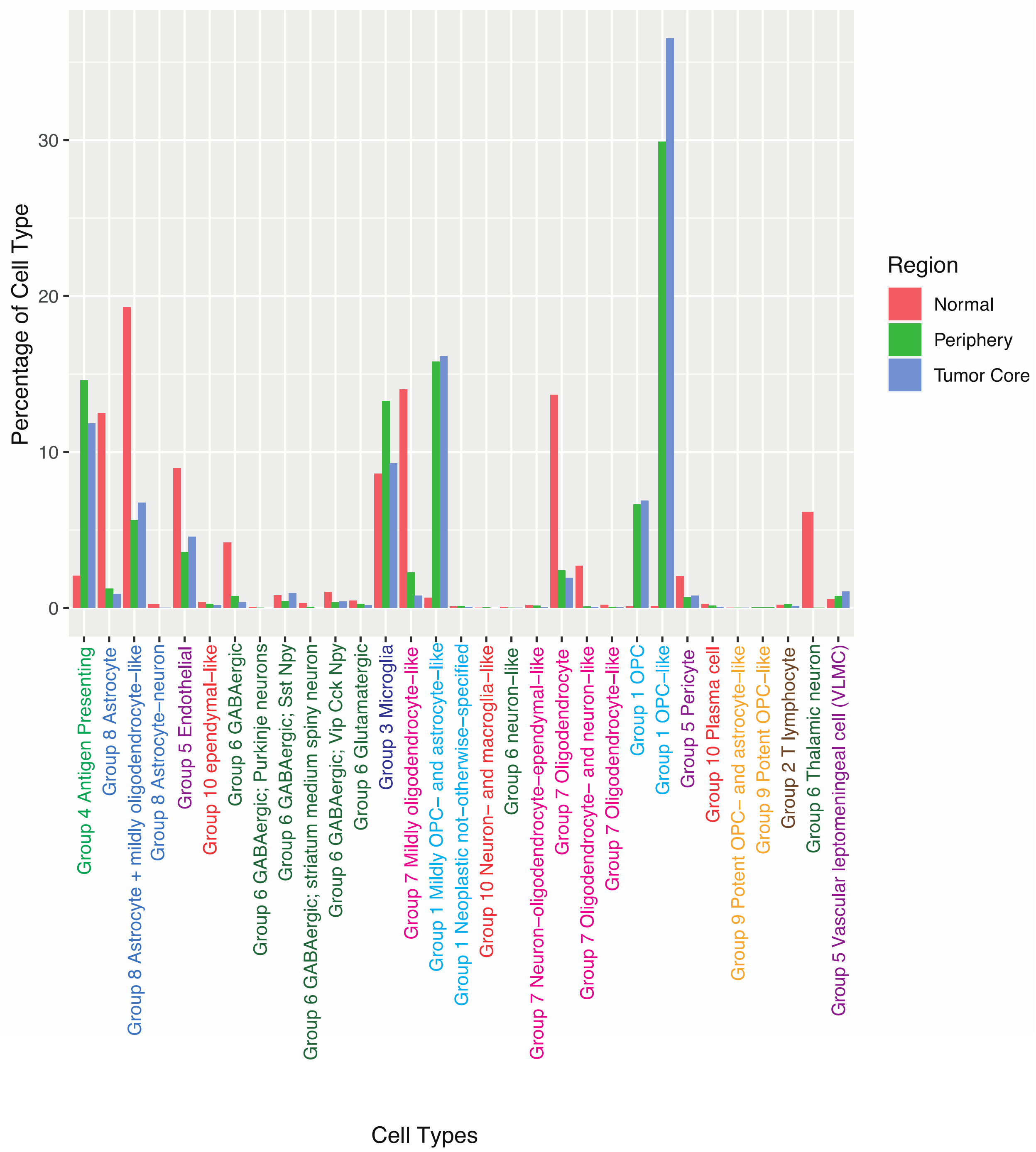


**Figure S3 –Collapsed individual cell clustering into 10 archetypal cell types.**
Tumor core, periphery, and normal tissues were delineated as shown in Figure 3B. Bar graph shows percentage of cells of each type within each region. Grouping of UMAP clusters into 10 archetypal cell types is indicated. Cells are from unirradiated tumor-bearing nPHA$^{FL/FL}$ mouse brain shown in Figure 3B. All 4 samples were grouped utilizing the above described 10 archetypal cell types.

| Name | Ensembl ID | Annotation |
|---|---|---|
| Acsbg1 | ENSMUSG00000032281 | Astrocytes |
| Aqp4 | ENSMUSG00000024411 | Astrocytes |
| Cdh20 | ENSMUSG00000050840 | Astrocytes |
| Clmn | ENSMUSG00000021097 | Astrocytes |
| Gfap | ENSMUSG00000020932 | Astrocytes |
| Gli3 | ENSMUSG00000021318 | Astrocytes |
| Id2 | ENSMUSG00000020644 | Astrocytes |
| Mapk4 | ENSMUSG00000024558 | Astrocytes |
| Ntsr2 | ENSMUSG00000020591 | Astrocytes |
| Pde7b | ENSMUSG00000019990 | Astrocytes |
| Rfx4 | ENSMUSG00000020037 | Astrocytes |
| Rorb | ENSMUSG00000036192 | Astrocytes |
| Slc39a12 | ENSMUSG00000036949 | Astrocytes |
| Arhgap12 | ENSMUSG00000041225 | CA1-ProS |
| Fibcd1 | ENSMUSG00000026841 | CA1-ProS |

| | | |
|---|---|---|
| Sipa1l3 | ENSMUSG00000030583 | CA1-ProS |
| Wfs1 | ENSMUSG00000039474 | CA1-ProS |
| 2010300C02Rik | ENSMUSG00000026090 | CA2 |
| Arhgef28 | ENSMUSG00000021662 | CA2 |
| Bcl11b | ENSMUSG00000048251 | CA2 |
| Bhlhe22 | ENSMUSG00000025128 | CA2 |
| Cabp7 | ENSMUSG00000009075 | CA2 |
| Cpne4 | ENSMUSG00000032564 | CA2 |
| Igfbp4 | ENSMUSG00000017493 | CA2 |
| Necab2 | ENSMUSG00000031837 | CA2 |
| Prdm8 | ENSMUSG00000035456 | CA2 |
| Strip2 | ENSMUSG00000039629 | CA2 |
| Syndig1 | ENSMUSG00000074736 | CA2 |
| Cpne6 | ENSMUSG00000022212 | CA3 |
| Epha4 | ENSMUSG00000026235 | CA3 |

| | | |
|---|---|---|
| Hat1 | ENSMUSG00000027018 | CA3 |
| Neurod6 | ENSMUSG00000037984 | CA3 |
| Npy2r | ENSMUSG00000028004 | CA3 |
| Nrp2 | ENSMUSG00000025969 | CA3 |
| Shisa6 | ENSMUSG00000053930 | CA3 |
| Adamtsl1 | ENSMUSG00000066113 | CT SUB |
| Prss35 | ENSMUSG00000033491 | CT SUB |
| Rmst | ENSMUSG00000112117 | CT SUB |
| Zfpm2 | ENSMUSG00000022306 | CT SUB |
| Cacna2d2 | ENSMUSG00000010066 | Cajal-Retzius cells |
| Calb2 | ENSMUSG00000003657 | Cajal-Retzius cells |
| Cd24a | ENSMUSG00000047139 | Cajal-Retzius cells |
| Cdh4 | ENSMUSG00000000305 | Cajal-Retzius cells |
| Ebf3 | ENSMUSG00000010476 | Cajal-Retzius cells |
| Kctd8 | ENSMUSG00000037653 | Cajal-Retzius cells |

| | | |
|---|---|---|
| Pdzd2 | ENSMUSG00000022197 | Cajal-Retzius cells |
| Slc17a6 | ENSMUSG00000030500 | Cajal-Retzius cells |
| Tmem163 | ENSMUSG00000026347 | Cajal-Retzius cells |
| Trp73 | ENSMUSG00000029026 | Cajal-Retzius cells |
| Bdnf | ENSMUSG00000048482 | Car3 |
| Cux2 | ENSMUSG00000042589 | Car3 |
| Mdga1 | ENSMUSG00000043557 | Car3 |
| Tmem132d | ENSMUSG00000034310 | Car3 |
| Cdh9 | ENSMUSG00000025370 | Dentate gyrus granule cells |
| Orai2 | ENSMUSG00000039747 | Dentate gyrus granule cells |
| Prox1 | ENSMUSG00000010175 | Dentate gyrus granule cells |
| Rasl10a | ENSMUSG00000034209 | Dentate gyrus granule cells |
| Tanc1 | ENSMUSG00000035168 | Dentate gyrus granule cells |
| Acvrl1 | ENSMUSG00000000530 | Endothelial cells |

| | | |
|---|---|---|
| Adgrl4 | ENSMUSG00000039167 | Endothelial cells |
| Car4 | ENSMUSG00000000805 | Endothelial cells |
| Cd93 | ENSMUSG00000027435 | Endothelial cells |
| Cldn5 | ENSMUSG00000041378 | Endothelial cells |
| Cobll1 | ENSMUSG00000034903 | Endothelial cells |
| Emcn | ENSMUSG00000054690 | Endothelial cells |
| Fgd5 | ENSMUSG00000034037 | Endothelial cells |
| Fn1 | ENSMUSG00000026193 | Endothelial cells |
| Kdr | ENSMUSG00000062960 | Endothelial cells |
| Ly6a | ENSMUSG00000075602 | Endothelial cells |
| Mecom | ENSMUSG00000027684 | Endothelial cells |
| Nostrin | ENSMUSG00000034738 | Endothelial cells |
| Paqr5 | ENSMUSG00000032278 | Endothelial cells |
| Pecam1 | ENSMUSG00000020717 | Endothelial cells |
| Pglyrp1 | ENSMUSG00000030413 | Endothelial cells |
| Slfn5 | ENSMUSG00000054404 | Endothelial cells |
| Sox17 | ENSMUSG00000025902 | Endothelial cells |
| Zfp366 | ENSMUSG00000050919 | Endothelial cells |
| Dpyd | ENSMUSG00000033308 | L2 IT ENTl |
| Nrn1 | ENSMUSG00000039114 | L2 IT ENTl |
| Pkib | ENSMUSG00000019876 | L2 IT ENTl |
| Plcxd3 | ENSMUSG00000049148 | L2 IT ENTl |
| Sema3a | ENSMUSG00000028883 | L2 IT ENTl |
| Slc17a7 | ENSMUSG00000070570 | L2 IT ENTl |
| Adamts2 | ENSMUSG00000036545 | L2 IT ENTm |
| Cbln4 | ENSMUSG00000067578 | L2 IT ENTm |
| Cdh13 | ENSMUSG00000031841 | L2 IT ENTm |
| Gsg1l | ENSMUSG00000046182 | L2 IT ENTm |
| Nwd2 | ENSMUSG00000090061 | L2 IT ENTm |
| Rspo2 | ENSMUSG00000051920 | L2 IT ENTm |
| Unc13c | ENSMUSG00000062151 | L2 IT ENTm |
| Igfbp6 | ENSMUSG00000023046 | L2/3 IT CTX |
| Fign | ENSMUSG00000075324 | L2/3 IT ENTl |
| Cntnap5b | ENSMUSG00000067028 | L2/3 IT RHP |
| Tmem255a | ENSMUSG00000036502 | L2/3 IT RHP |
| Cd44 | ENSMUSG00000005087 | L3 IT ENT |
| Hpcal1 | ENSMUSG00000071379 | L3 IT ENT |
| Igfbp5 | ENSMUSG00000026185 | L3 IT ENT |
| Plch1 | ENSMUSG00000036834 | L3 IT ENT |

| | | |
|---|---|---|
| Cbln1 | ENSMUSG00000031654 | L4 RSP-ACA |
| Nell1 | ENSMUSG00000055409 | L4 RSP-ACA |
| Nrep | ENSMUSG00000042834 | L4 RSP-ACA |
| Opn3 | ENSMUSG00000026525 | L4 RSP-ACA |
| Parm1 | ENSMUSG00000034981 | L4 RSP-ACA |
| Pdzrn3 | ENSMUSG00000035357 | L4 RSP-ACA |
| Prph | ENSMUSG00000023484 | L4 RSP-ACA |
| Rspo1 | ENSMUSG00000028871 | L4 RSP-ACA |
| Arc | ENSMUSG00000022602 | L4/5 IT CTX |
| Fhod3 | ENSMUSG00000034295 | L4/5 IT CTX |
| Gfra2 | ENSMUSG00000022103 | L4/5 IT CTX |
| Kcnh5 | ENSMUSG00000034402 | L4/5 IT CTX |
| Rims3 | ENSMUSG00000032890 | L4/5 IT CTX |
| Deptor | ENSMUSG00000022419 | L5 IT CTX |
| Dkk3 | ENSMUSG00000030772 | L5 IT CTX |
| Hs3st2 | ENSMUSG00000046321 | L5 IT CTX |
| Fezf2 | ENSMUSG00000021743 | L5 NP CTX |
| Myl4 | ENSMUSG00000061086 | L5 NP CTX |
| Neto2 | ENSMUSG00000036902 | L5 NP CTX |
| Plcxd2 | ENSMUSG00000087141 | L5 NP CTX |
| Prr16 | ENSMUSG00000073565 | L5 NP CTX |
| Rnf152 | ENSMUSG00000047496 | L5 NP CTX |
| Angpt1 | ENSMUSG00000022309 | L5 PPP |
| Dpy19l1 | ENSMUSG00000043067 | L5 PPP |
| Ndst3 | ENSMUSG00000027977 | L5 PPP |
| Nts | ENSMUSG00000019890 | L5 PPP |
| Pou3f1 | ENSMUSG00000090125 | L5 PPP |
| Rxfp1 | ENSMUSG00000034009 | L5 PPP |
| Syt2 | ENSMUSG00000026452 | L5 PPP |
| Vat1l | ENSMUSG00000046844 | L5 PPP |
| Gm19410 | ENSMUSG00000109372 | L5 PT CTX |
| Npnt | ENSMUSG00000040998 | L5 PT CTX |
| Foxp2 | ENSMUSG00000029563 | L6 CT CTX |
| Gadd45a | ENSMUSG00000036390 | L6 CT CTX |
| Garnl3 | ENSMUSG00000038860 | L6 CT CTX |
| Ppp1r1b | ENSMUSG00000061718 | L6 CT CTX |
| Rprm | ENSMUSG00000075334 | L6 CT CTX |
| Trbc2 | ENSMUSG00000076498 | L6 CT CTX |
| Galnt14 | ENSMUSG00000024064 | L6 IT ENTl |
| Spag16 | ENSMUSG00000053153 | L6 IT ENTl |
| Syt17 | ENSMUSG00000058420 | L6 IT ENTl |
| Ccn2 | ENSMUSG00000019997 | L6b CTX |

| | | |
|---|---|---|
| Cplx3 | ENSMUSG00000039714 | L6b CTX |
| Gng12 | ENSMUSG00000036402 | L6b CTX |
| Sdk2 | ENSMUSG00000041592 | L6b CTX |
| Cwh43 | ENSMUSG00000029154 | L6b/CT ENT |
| Igsf21 | ENSMUSG00000040972 | L6b/CT ENT |
| Sox11 | ENSMUSG00000063632 | L6b/CT ENT |
| Dner | ENSMUSG00000036766 | Lamp5 interneurons |
| Gad1 | ENSMUSG00000070880 | Lamp5 interneurons |
| Gad2 | ENSMUSG00000026787 | Lamp5 interneurons |
| Hapln1 | ENSMUSG00000021613 | Lamp5 interneurons |
| Lamp5 | ENSMUSG00000027270 | Lamp5 interneurons |
| Pde11a | ENSMUSG00000075270 | Lamp5 interneurons |
| Rasgrf2 | ENSMUSG00000021708 | Lamp5 interneurons |
| Cpne8 | ENSMUSG00000052560 | Meis2 |
| Grik3 | ENSMUSG00000001985 | Meis2 |
| Htr1f | ENSMUSG00000050783 | Meis2 |
| Meis2 | ENSMUSG00000027210 | Meis2 |
| Myo16 | ENSMUSG00000039057 | Meis2 |
| Sema3e | ENSMUSG00000063531 | Meis2 |
| Slc6a3 | ENSMUSG00000021609 | Meis2 |
| Syt6 | ENSMUSG00000027849 | Meis2 |
| Th | ENSMUSG00000000214 | Meis2 |
| Arhgap25 | ENSMUSG00000030047 | Microglia / perivascular macrophages |
| Cd300c2 | ENSMUSG00000044811 | Microglia / perivascular macrophages |
| Cd53 | ENSMUSG00000040747 | Microglia / perivascular macrophages |
| Cd68 | ENSMUSG00000018774 | Microglia / perivascular macrophages |
| Ikzf1 | ENSMUSG00000018654 | Microglia / perivascular macrophages |
| Laptm5 | ENSMUSG00000028581 | Microglia / perivascular macrophages |
| Lyz2 | ENSMUSG00000069516 | Microglia / perivascular macrophages |
| Siglech | ENSMUSG00000051504 | Microglia / perivascular macrophages |
| Sla | ENSMUSG00000022372 | Microglia / perivascular macrophages |
| Spi1 | ENSMUSG00000002111 | Microglia / perivascular macrophages |
| Trem2 | ENSMUSG00000023992 | Microglia / perivascular macrophages |

| | | |
|---|---|---|
| Pcsk5 | ENSMUSG00000024713 | NP PPP |
| Satb2 | ENSMUSG00000038331 | NP PPP |
| Tle4 | ENSMUSG00000024642 | NP PPP |
| Tox | ENSMUSG00000041272 | NP PPP |
| Cntn6 | ENSMUSG00000030092 | NP SUB |
| Nxph3 | ENSMUSG00000046719 | NP SUB |
| Sema5b | ENSMUSG00000052133 | NP SUB |
| Stard5 | ENSMUSG00000046027 | NP SUB |
| Vwc2l | ENSMUSG00000045648 | NP SUB |
| Gjc3 | ENSMUSG00000056966 | Oligodendrocytes |
| Gpr17 | ENSMUSG00000052229 | Oligodendrocytes |
| Opalin | ENSMUSG00000050121 | Oligodendrocytes |
| Sema3d | ENSMUSG00000040254 | Oligodendrocytes |
| Sema6a | ENSMUSG00000019647 | Oligodendrocytes |
| Sox10 | ENSMUSG00000033006 | Oligodendrocytes |
| Zfp536 | ENSMUSG00000043456 | Oligodendrocytes |
| Acta2 | ENSMUSG00000035783 | Pericytes / smooth muscle cells |
| Ano1 | ENSMUSG00000031075 | Pericytes / smooth muscle cells |
| Arhgap6 | ENSMUSG00000031355 | Pericytes / smooth muscle cells |
| Carmn | ENSMUSG00000097324 | Pericytes / smooth muscle cells |
| Cspg4 | ENSMUSG00000032911 | Pericytes / smooth muscle cells |
| Fos | ENSMUSG00000021250 | Pericytes / smooth muscle cells |
| Gucy1a1 | ENSMUSG00000033910 | Pericytes / smooth muscle cells |
| Inpp4b | ENSMUSG00000037940 | Pericytes / smooth muscle cells |
| Nr2f2 | ENSMUSG00000030551 | Pericytes / smooth muscle cells |
| Pip5k1b | ENSMUSG00000024867 | Pericytes / smooth muscle cells |
| Plekha2 | ENSMUSG00000031557 | Pericytes / smooth muscle cells |
| Pln | ENSMUSG00000038583 | Pericytes / smooth muscle cells |
| Sncg | ENSMUSG00000023064 | Pericytes / smooth muscle cells |
| Sntb1 | ENSMUSG00000060429 | Pericytes / smooth muscle cells |
| Btbd11 | ENSMUSG00000020042 | Pvalb interneurons |

| | | |
|---|---|---|
| Cntnap4 | ENSMUSG00000031772 | Pvalb interneurons |
| Eya4 | ENSMUSG00000010461 | Pvalb interneurons |
| Kcnmb2 | ENSMUSG00000037610 | Pvalb interneurons |
| Pvalb | ENSMUSG00000005716 | Pvalb interneurons |
| Slit2 | ENSMUSG00000031558 | Pvalb interneurons |
| Bhlhe40 | ENSMUSG00000030103 | SUB-ProS |
| Cdh6 | ENSMUSG00000039385 | SUB-ProS |
| Gm2115 | ENSMUSG00000097789 | SUB-ProS |
| Trpc4 | ENSMUSG00000027748 | SUB-ProS |
| Col19a1 | ENSMUSG00000026141 | Sncg interneurons |
| Kctd12 | ENSMUSG00000098557 | Sncg interneurons |
| Necab1 | ENSMUSG00000040536 | Sncg interneurons |
| Slc44a5 | ENSMUSG00000028360 | Sncg interneurons |
| Chodl | ENSMUSG00000022860 | Sst Chodl inteneurons |
| Chrm2 | ENSMUSG00000045613 | Sst Chodl inteneurons |
| Cort | ENSMUSG00000028971 | Sst Chodl inteneurons |
| Ndst4 | ENSMUSG00000027971 | Sst Chodl inteneurons |
| Sst | ENSMUSG00000004366 | Sst Chodl inteneurons |
| Tacr1 | ENSMUSG00000030043 | Sst Chodl inteneurons |
| Calb1 | ENSMUSG00000028222 | Sst interneurons |
| Lypd6 | ENSMUSG00000050447 | Sst interneurons |
| Pdyn | ENSMUSG00000027400 | Sst interneurons |
| Rab3b | ENSMUSG00000003411 | Sst interneurons |
| Rbp4 | ENSMUSG00000024990 | Sst interneurons |
| Aldh1a2 | ENSMUSG00000013584 | VLMC |
| Col1a1 | ENSMUSG00000001506 | VLMC |
| Col6a1 | ENSMUSG00000001119 | VLMC |
| Cyp1b1 | ENSMUSG00000024087 | VLMC |
| Dcn | ENSMUSG00000019929 | VLMC |
| Fmod | ENSMUSG00000041559 | VLMC |
| Gjb2 | ENSMUSG00000046352 | VLMC |
| Igf2 | ENSMUSG00000048583 | VLMC |
| Pdgfra | ENSMUSG00000029231 | VLMC |
| Ror1 | ENSMUSG00000035305 | VLMC |
| Slc13a4 | ENSMUSG00000029843 | VLMC |
| Spp1 | ENSMUSG00000029304 | VLMC |
| Chat | ENSMUSG00000021919 | Vip interneurons |
| Crh | ENSMUSG00000049796 | Vip interneurons |
| Igf1 | ENSMUSG00000020053 | Vip interneurons |
| Penk | ENSMUSG00000045573 | Vip interneurons |
| Pthlh | ENSMUSG00000048776 | Vip interneurons |
| Sorcs3 | ENSMUSG00000063434 | Vip interneurons |

| | | |
|---|---|---|
| Thsd7a | ENSMUSG00000032625 | Vip interneurons |
| Vip | ENSMUSG00000019772 | Vip interneurons |
| Acvr1 | ENSMUSG00000026836 | DMG Canonical Markers |
| Aldh1l1 | ENSMUSG00000030088 | DMG Canonical Markers |
| Atm | ENSMUSG00000034218 | DMG Canonical Markers |
| C1qc | ENSMUSG00000036896 | DMG Canonical Markers |
| Cd163 | ENSMUSG00000008845 | Macrophage |
| Cd34 | ENSMUSG00000016494 | DMG Canonical Markers |
| Cd3d | ENSMUSG00000032094 | T-lymphocyte |
| Cd4 | ENSMUSG00000023274 | T-lymphocyte |
| Cd8a | ENSMUSG00000053977 | T-lymphocyte |
| Cdkn1a | ENSMUSG00000023067 | DMG Canonical Markers |
| Cnp | ENSMUSG00000006782 | DMG Canonical Markers |
| Col1a2 | ENSMUSG00000029661 | DMG Canonical Markers |
| Fdft1 | ENSMUSG00000021273 | DMG Canonical Markers |
| Foxp3 | ENSMUSG00000039521 | DMG Canonical Markers |
| Gzma | ENSMUSG00000023132 | DMG Canonical Markers |
| H3f3a | ENSMUSG00000060743 | DMG Canonical Markers |
| Hmgcs1 | ENSMUSG00000093930 | DMG Canonical Markers |
| Il17a | ENSMUSG00000025929 | DMG Canonical Markers |
| Itgam | ENSMUSG00000030786 | DMG Canonical Markers |
| Gli1 | ENSMUSG00000025407 | DMG Canonical Markers |
| Gli2 | ENSMUSG00000048402 | DMG Canonical Markers |
| Mcm2 | ENSMUSG00000002870 | DMG Canonical Markers |
| Mdm2 | ENSMUSG00000002870 | DMG Canonical Markers |
| Mobp | ENSMUSG00000032517 | DMG Canonical Markers |
| Ms4a1 | ENSMUSG00000024673 | DMG Canonical Markers |
| Msr1 | ENSMUSG00000025044 | DMG Canonical Markers |
| Nrgn | ENSMUSG00000053310 | DMG Canonical Markers |
| Olig1 | ENSMUSG00000046160 | DMG Canonical Markers |
| Olig2 | ENSMUSG00000039830 | DMG Canonical Markers |
| P2ry12 | ENSMUSG00000036353 | DMG Canonical Markers |

| | | |
|---|---|---|
| Pax6 | ENSMUSG00000027168 | DMG Canonical Markers |
| Pik3ca | ENSMUSG00000027665 | DMG Canonical Markers |
| Ppm1d | ENSMUSG00000020525 | DMG Canonical Markers |
| Prf1 | ENSMUSG00000037202 | DMG Canonical Markers |
| Ptch1 | ENSMUSG00000021466 | DMG Canonical Markers |
| Pten | ENSMUSG00000013663 | DMG Canonical Markers |
| Ptprc | ENSMUSG00000026395 | DMG Canonical Markers |
| Rbfox3 | ENSMUSG00000025576 | DMG Canonical Markers |
| Slc1a2 | ENSMUSG00000005089 | DMG Canonical Markers |
| Slc32a1 | ENSMUSG00000037771 | DMG Canonical Markers |
| Sox2 | ENSMUSG00000074637 | DMG Canonical Markers |
| Sox4 | ENSMUSG00000076431 | DMG Canonical Markers |
| Sox8 | ENSMUSG00000024176 | DMG Canonical Markers |
| Sox9 | ENSMUSG00000000567 | DMG Canonical Markers |
| Tmem144 | ENSMUSG00000027956 | DMG Canonical Markers |
| Tmem173 | ENSMUSG00000024349 | DMG Canonical Markers |
| Top2a | ENSMUSG00000020914 | DMG Canonical Markers |
| Trp53 | ENSMUSG00000059552 | DMG Canonical Markers |
| Tyrobp | ENSMUSG00000030579 | DMG Canonical Markers |
| Vwf | ENSMUSG00000001930 | DMG Canonical Markers |

Table S2: Collapsed cell clustering

| Cell Type Identified by Xenium platform | Collapsed Cell Clustering |
| --- | --- |
| Mildly OPC- and astrocyte-like<br>Neoplastic not-otherwise specified<br>OPC<br>OPC-like | Neoplastic (1) |
| T-lymphocyte | T-Lymphocyte Cell (2) |
| Microglia | Microglia (3) |
| Antigen Presenting Cell | Antigen Presenting Cell (4) |
| Endothelial | Endothelium (5) |
| GABAergic<br>Purkinje Neurons<br>GABAergic; Sst Npy<br>GABAergic; stratum medium spiny neuron<br>GABAergic; Vip Cck Npy<br>Glutamatergic<br>Neuron-like<br>Thalamic Neuron | Neuron (6) |
| Mildly Oligodendrocyte-like<br>Neuron-oligodendrocyte-ependymal-like<br>Oligodendrocyte<br>Oligodendrocyte-and neuron-like<br>Oligodendrocyte-like | Normal Oligodendrocyte (7) |
| Astrocyte<br>Astrocyte + mildly oligodendrocyte like<br>Astrocyte neuron | Normal Astrocyte (8) |
| Potent OPC-and Astrocyte like<br>Potent OPC-like | Normal Oligodendrocyte precursor cell (9) |
| Ependymal – like<br>Plasma Cell | Other: Ependymal cells, Plasma cell etc. (10) |

**Table S3**

Atm$^{FL/+}$ with and without irradiation

| Differentially Expressed Genes | p_val | avg_log2FC | pct.1 | pct.2 | p_val_adj |
|---|---|---|---|---|---|
| Cd24a | 0 | 0.45465633 | 0.535 | 0.371 | 0 |
| Cdkn1a | 0 | 0.82625756 | 0.305 | 0.156 | 0 |
| Cntn6 | 0 | -0.9533859 | 0.074 | 0.217 | 0 |
| Col19a1 | 0 | -0.6388915 | 0.414 | 0.614 | 0 |
| Cpne8 | 0 | 0.65663785 | 0.701 | 0.535 | 0 |
| Dpy19l1 | 0 | -0.6679676 | 0.715 | 0.862 | 0 |
| Gjc3 | 0 | 0.38412926 | 0.966 | 0.939 | 0 |
| Gng12 | 0 | -0.2903972 | 0.957 | 0.978 | 0 |
| Gpr17 | 0 | -1.0042095 | 0.675 | 0.903 | 0 |
| Igfbp4 | 0 | 0.84272815 | 0.599 | 0.399 | 0 |
| Mobp | 0 | -1.0655137 | 0.061 | 0.181 | 0 |
| Neto2 | 0 | 0.59804994 | 0.75 | 0.616 | 0 |
| Ntsr2 | 0 | -0.8856547 | 0.11 | 0.251 | 0 |
| Pde7b | 0 | 0.64834639 | 0.482 | 0.33 | 0 |
| Pou3f1 | 0 | -1.7321007 | 0.431 | 0.863 | 0 |
| Pten | 0 | 0.43419965 | 0.942 | 0.887 | 0 |
| Rprm | 0 | -0.5341461 | 0.458 | 0.61 | 0 |
| Sema3a | 0 | 0.81370153 | 0.34 | 0.185 | 0 |
| Sema3d | 0 | 1.13596382 | 0.823 | 0.553 | 0 |
| Sox2 | 0 | -0.330949 | 0.967 | 0.986 | 0 |
| Sox8 | 0 | -0.5154122 | 0.781 | 0.894 | 0 |
| Sox9 | 0 | -1.1872076 | 0.186 | 0.442 | 0 |
| Syt6 | 0 | 0.86444409 | 0.279 | 0.125 | 0 |
| Meis2 | 1.25E-283 | 0.5602549 | 0.544 | 0.408 | 3.73E-281 |
| Zfp536 | 1.47E-271 | 0.61154868 | 0.413 | 0.271 | 4.39E-269 |
| Arc | 1.68E-271 | 0.88155626 | 0.209 | 0.095 | 5.00E-269 |
| Tmem132d | 1.05E-232 | -0.4974553 | 0.368 | 0.496 | 3.13E-230 |
| Dner | 1.17E-226 | -0.3220224 | 0.756 | 0.838 | 3.50E-224 |
| Rab3b | 4.26E-224 | 0.53254171 | 0.462 | 0.335 | 1.27E-221 |
| Fdft1 | 3.24E-220 | -0.4163337 | 0.486 | 0.608 | 9.65E-218 |
| Cnp | 1.80E-209 | 0.30838304 | 0.78 | 0.692 | 5.38E-207 |
| Aldh1l1 | 7.73E-199 | 0.8521914 | 0.177 | 0.086 | 2.30E-196 |
| Mcm2 | 1.32E-165 | 0.41357323 | 0.621 | 0.53 | 3.94E-163 |
| Tox | 9.79E-161 | -0.4969958 | 0.158 | 0.25 | 2.92E-158 |

| | | | | | |
|---|---|---|---|---|---|
| Top2a | 1.10E-159 | 0.28204286 | 0.889 | 0.836 | 3.28E-157 |
| Arhgap25 | 5.41E-144 | -0.5362523 | 0.055 | 0.114 | 1.61E-141 |
| Gucy1a1 | 2.97E-139 | 0.51777822 | 0.239 | 0.151 | 8.85E-137 |
| Rorb | 8.12E-131 | 0.40191759 | 0.5 | 0.407 | 2.42E-128 |
| Cdh13 | 1.10E-125 | 0.33294844 | 0.627 | 0.547 | 3.28E-123 |
| Fos | 3.16E-123 | 0.57973269 | 0.237 | 0.156 | 9.43E-121 |
| Gfap | 7.85E-122 | -0.3460217 | 0.283 | 0.381 | 2.34E-119 |
| Nrep | 5.20E-121 | -0.2983228 | 0.502 | 0.596 | 1.55E-118 |
| Thsd7a | 3.60E-119 | 0.44123699 | 0.373 | 0.284 | 1.07E-116 |
| 2010300C02Rik | 1.62E-111 | -0.4515898 | 0.085 | 0.145 | 4.83E-109 |
| Lyz2 | 1.82E-111 | 0.49140701 | 0.206 | 0.13 | 5.42E-109 |
| Dkk3 | 1.08E-109 | 0.46464082 | 0.231 | 0.153 | 3.21E-107 |
| Plekha2 | 4.67E-104 | 0.29636309 | 0.629 | 0.554 | 1.39E-101 |
| Tanc1 | 2.84E-101 | 0.27519069 | 0.663 | 0.592 | 8.45E-99 |
| Calb2 | 1.90E-94 | -0.9437316 | 0.06 | 0.107 | 5.65E-92 |
| Plcxd2 | 2.53E-88 | -0.4390091 | 0.07 | 0.119 | 7.55E-86 |
| Prss35 | 4.92E-85 | 0.48382686 | 0.12 | 0.069 | 1.47E-82 |
| Kcnh5 | 1.69E-78 | 0.36233524 | 0.357 | 0.285 | 5.02E-76 |
| Arhgef28 | 2.13E-78 | 0.40198553 | 0.134 | 0.081 | 6.34E-76 |
| Kctd12 | 8.34E-77 | 0.31845716 | 0.372 | 0.298 | 2.49E-74 |
| Rims3 | 6.20E-76 | -0.4395223 | 0.105 | 0.158 | 1.85E-73 |
| Pdzrn3 | 2.95E-74 | 0.49602445 | 0.136 | 0.085 | 8.78E-72 |
| Prdm8 | 3.32E-74 | -0.3600763 | 0.082 | 0.131 | 9.90E-72 |
| Lypd6 | 1.63E-63 | -0.3019273 | 0.289 | 0.354 | 4.85E-61 |
| Igsf21 | 9.68E-63 | 0.28647183 | 0.395 | 0.33 | 2.88E-60 |
| Hpcal1 | 1.41E-60 | -0.3192566 | 0.273 | 0.336 | 4.19E-58 |
| Fn1 | 6.35E-54 | 0.30947881 | 0.215 | 0.161 | 1.89E-51 |
| Foxp2 | 2.29E-51 | -0.2855498 | 0.201 | 0.255 | 6.82E-49 |
| Bcl11b | 7.91E-49 | 0.32651332 | 0.103 | 0.066 | 2.36E-46 |
| Plch1 | 2.90E-48 | 0.31871345 | 0.129 | 0.088 | 8.66E-46 |
| Laptm5 | 1.40E-43 | -0.3700916 | 0.094 | 0.131 | 4.18E-41 |
| Zfpm2 | 4.54E-39 | 0.26389695 | 0.249 | 0.203 | 1.35E-36 |
| Cabp7 | 4.65E-39 | 0.30337275 | 0.131 | 0.094 | 1.38E-36 |
| Igfbp5 | 1.37E-33 | 0.38005264 | 0.195 | 0.156 | 4.07E-31 |
| C1qc | 2.01E-33 | -0.3428421 | 0.138 | 0.174 | 5.99E-31 |

Table S4

Atm$^{FL/FL}$ with and without irradiation DEGs

| Differentially Expressed Genes | P-value | Average $Log_2$fold change | pct.1 | pct.2 | p-value adjusted |
|---|---|---|---|---|---|
| Ly6a | 0 | 2.98215354 | 0.859 | 0.172 | 0 |
| Lyz2 | 0 | 4.23344401 | 0.919 | 0.107 | 0 |
| Igf1 | 3.94E-222 | 1.80694116 | 0.267 | 0.019 | 1.18E-219 |
| Ndst3 | 1.30E-199 | 1.85249795 | 0.509 | 0.104 | 3.87E-197 |
| Cd44 | 2.51E-197 | 2.13172941 | 0.407 | 0.065 | 7.49E-195 |
| Acsbg1 | 5.78E-183 | 1.43591491 | 0.946 | 0.618 | 1.72E-180 |
| Pde11a | 1.97E-154 | 1.48705741 | 0.196 | 0.015 | 5.88E-152 |
| Slfn5 | 1.22E-144 | 1.60425171 | 0.284 | 0.041 | 3.64E-142 |
| Hmgcs1 | 2.43E-132 | -1.5009082 | 0.589 | 0.857 | 7.24E-130 |
| C1qc | 5.77E-113 | 1.57155156 | 0.65 | 0.275 | 1.72E-110 |
| Ccn2 | 2.64E-110 | 1.5840533 | 0.208 | 0.028 | 7.88E-108 |
| Plch1 | 4.18E-108 | 1.98761893 | 0.314 | 0.07 | 1.25E-105 |
| H3f3a | 2.70E-102 | -0.482292 | 0.996 | 0.999 | 8.04E-100 |
| Ptprc | 1.74E-95 | 1.5100652 | 0.179 | 0.024 | 5.17E-93 |
| Laptm5 | 1.08E-93 | 1.41268304 | 0.489 | 0.173 | 3.23E-91 |
| Pdgfra | 6.05E-92 | -0.4518301 | 0.99 | 1 | 1.80E-89 |
| Cd68 | 4.20E-78 | 1.4727131 | 0.341 | 0.104 | 1.25E-75 |
| Cspg4 | 5.05E-78 | -0.6589742 | 0.898 | 0.976 | 1.51E-75 |
| Fdft1 | 1.09E-75 | -1.4165217 | 0.347 | 0.66 | 3.25E-73 |
| Olig2 | 8.61E-65 | -0.4715824 | 0.976 | 0.997 | 2.56E-62 |
| Top2a | 4.92E-64 | -1.7486486 | 0.186 | 0.521 | 1.47E-61 |
| Tyrobp | 7.61E-63 | 1.18999988 | 0.336 | 0.114 | 2.27E-60 |
| Cdh4 | 1.30E-60 | -1.4784587 | 0.181 | 0.496 | 3.86E-58 |
| Pten | 1.57E-58 | 0.5696656 | 0.967 | 0.879 | 4.68E-56 |
| Cd53 | 2.70E-56 | 1.19715954 | 0.243 | 0.07 | 8.04E-54 |
| Cabp7 | 5.69E-55 | 1.08922467 | 0.346 | 0.13 | 1.69E-52 |
| Gng12 | 6.32E-54 | -0.5100133 | 0.945 | 0.975 | 1.88E-51 |
| Tmem132d | 7.66E-49 | -0.9016209 | 0.501 | 0.724 | 2.28E-46 |
| Cd24a | 1.40E-45 | 1.06924757 | 0.528 | 0.295 | 4.17E-43 |
| Arhgef28 | 1.35E-44 | 0.96203677 | 0.347 | 0.146 | 4.02E-42 |
| Col1a2 | 6.15E-44 | 1.23832124 | 0.141 | 0.032 | 1.83E-41 |
| Sdk2 | 8.38E-43 | 0.88307863 | 0.129 | 0.028 | 2.50E-40 |
| Sox2 | 1.27E-42 | -0.4453391 | 0.917 | 0.964 | 3.80E-40 |
| Pou3f1 | 6.24E-40 | 0.77351178 | 0.707 | 0.505 | 1.86E-37 |

| | | | | | |
|---|---|---|---|---|---|
| Slc1a2 | 1.22E-39 | 0.37697851 | 0.99 | 0.97 | 3.62E-37 |
| Col6a1 | 6.08E-38 | 1.09447317 | 0.101 | 0.02 | 1.81E-35 |
| Tmem144 | 8.59E-38 | 0.91615103 | 0.397 | 0.196 | 2.56E-35 |
| Parm1 | 2.67E-36 | 1.10065967 | 0.249 | 0.096 | 7.95E-34 |
| Slit2 | 2.97E-34 | -0.862754 | 0.306 | 0.542 | 8.86E-32 |
| Dner | 2.81E-33 | -0.3945525 | 0.903 | 0.945 | 8.38E-31 |
| Sox4 | 5.49E-33 | -0.5721428 | 0.69 | 0.82 | 1.63E-30 |
| Gfap | 1.25E-31 | 0.60138056 | 0.836 | 0.692 | 3.72E-29 |
| Fn1 | 1.56E-30 | 1.00256142 | 0.269 | 0.118 | 4.65E-28 |
| Meis2 | 2.67E-29 | -0.5883184 | 0.514 | 0.694 | 7.95E-27 |
| Igfbp4 | 1.20E-27 | 0.78252666 | 0.63 | 0.464 | 3.57E-25 |
| Gadd45a | 1.35E-27 | 0.95178348 | 0.364 | 0.195 | 4.01E-25 |
| Bhlhe40 | 7.47E-26 | 0.77620985 | 0.428 | 0.252 | 2.23E-23 |
| Angpt1 | 6.51E-24 | 0.76308213 | 0.312 | 0.163 | 1.94E-21 |
| Gm2115 | 6.56E-24 | 0.74811419 | 0.182 | 0.073 | 1.95E-21 |
| Cd300c2 | 7.19E-23 | 0.72751193 | 0.168 | 0.066 | 2.14E-20 |
| Lypd6 | 9.88E-23 | 0.77382067 | 0.35 | 0.198 | 2.94E-20 |
| Necab2 | 6.97E-22 | -0.7398512 | 0.334 | 0.503 | 2.08E-19 |
| Hpcal1 | 7.67E-22 | 0.61519381 | 0.454 | 0.284 | 2.29E-19 |
| Gucy1a1 | 8.65E-22 | -0.7476419 | 0.297 | 0.474 | 2.58E-19 |
| Pdzd2 | 9.60E-22 | 0.61992912 | 0.501 | 0.335 | 2.86E-19 |
| Trp53 | 2.11E-21 | 0.70218841 | 0.11 | 0.035 | 6.30E-19 |
| Gjc3 | 4.19E-21 | 0.29941251 | 0.99 | 0.986 | 1.25E-18 |
| Cobll1 | 1.26E-20 | 0.47969117 | 0.71 | 0.556 | 3.74E-18 |
| Acta2 | 1.89E-20 | 0.82247983 | 0.199 | 0.09 | 5.62E-18 |
| Aqp4 | 4.46E-20 | 0.74658106 | 0.448 | 0.292 | 1.33E-17 |
| Wfs1 | 1.35E-19 | 0.71863796 | 0.205 | 0.096 | 4.03E-17 |
| 2010300C02Rik | 2.00E-19 | -0.8845026 | 0.121 | 0.275 | 5.97E-17 |
| Cntnap5b | 3.56E-19 | -0.93947 | 0.084 | 0.229 | 1.06E-16 |
| Trem2 | 6.39E-19 | 0.66197924 | 0.193 | 0.089 | 1.90E-16 |
| Rab3b | 9.52E-19 | -0.6141688 | 0.31 | 0.477 | 2.84E-16 |
| Dpyd | 6.21E-18 | 0.51193471 | 0.596 | 0.449 | 1.85E-15 |
| Tox | 1.80E-17 | -0.6745374 | 0.193 | 0.35 | 5.35E-15 |
| Sema6a | 2.21E-17 | -0.4048593 | 0.777 | 0.844 | 6.59E-15 |
| Slc44a5 | 5.01E-16 | -0.7426964 | 0.165 | 0.31 | 1.49E-13 |
| Gfra2 | 5.07E-16 | 0.73275394 | 0.299 | 0.18 | 1.51E-13 |
| Slc39a12 | 1.45E-15 | 0.63998648 | 0.131 | 0.055 | 4.32E-13 |
| Spi1 | 3.26E-15 | 0.50595543 | 0.102 | 0.038 | 9.72E-13 |

Table S5: Top Cell:Ligand receptor

Atm intact

| | n_spots | n_spots_sig | n_spots_sig_pval | n_cci_sig_celltype | n-spot_cci_celltype | n-spot_cci_sig_celltype |
|---|---|---|---|---|---|---|
| Col1a2_Cd93 | 23723 | 2694 | 5124 | 14 | 14021 | 6686 |
| Col1a1_Cd93 1 | 10095 | 1855 | 4283 | 10 | 8528 | 5338 |
| Fn1_Cd44 | 16090 | 1839 | 3039 | 16 | 13541 | 7294 |
| Col1a2_Cd44 | 8273 | 759 | 1273 | 17 | 3847 | 2469 |
| Spp1_Cd44 | 6069 | 732 | 1269 | 14 | 4585 | 3433 |
| Col1a1_Cd44 | 3770 | 592 | 1128 | 14 | 2863 | 2024 |
| Sema3a_Nrp2 | 9926 | 501 | 1055 | 15 | 1840 | 442 |
| Nts_Ntsr2 | 5486 | 440 | 863 | 25 | 1845 | 678 |
| Ccn2_Itgam | 6480 | 437 | 893 | 10 | 1364 | 1169 |

Atm intact with irradiation

| | n_spots | n_spots_sig | n_spots_sig_pval | n_cci_sig_celltype | n-spot_cci_celltype | n-spot_cci_sig_celltype |
|---|---|---|---|---|---|---|
| Fn1_Cd44 | 40314 | 3450 | 6849 | 15 | 30215 | 22395 |
| Col1a2_Cd93 | 41079 | 2149 | 3909 | 9 | 18043 | 14757 |
| Col1a1_Cd93 | 21745 | 1729 | 3659 | 8 | 13885 | 11506 |
| Col1a2_Cd44 | 20598 | 1468 | 2270 | 10 | 13946 | 11471 |
| Spp1_Cd44 | 19217 | 1267 | 2254 | 13 | 12441 | 9660 |
| Col1a1_Cd44 | 11956 | 1139 | 1803 | 10 | 10470 | 8721 |
| Ccn2_Itgam | 16744 | 801 | 1542 | 10 | 3629 | 3134 |
| Sema3a_Nrp2 | 27072 | 669 | 2032 | 2 | 3381 | 1172 |
| Nts_Ntsr2 | 7012 | 629 | 1407 | 36 | 2629 | 1435 |

Atm null

| | n_spots | n_spots_sig | n_spots_sig_pval | n_cci_sig_celltype | n-spot_cci_celltype | n-spot_cci_sig_celltype |
|---|---|---|---|---|---|---|
| Fn1_Cd44 | 4784 | 783 | 1183 | 12 | 4999 | 4292 |
| Col1a2_Cd93 | 4426 | 447 | 736 | 8 | 3291 | 3123 |
| Col1a1_Cd93 | 2580 | 426 | 633 | 8 | 2987 | 2811 |
| Nts_Ntsr2 | 2199 | 326 | 445 | 34 | 2958 | 1776 |
| Spp1_Cd44 | 2371 | 289 | 489 | 10 | 1738 | 1402 |
| Col1a2_Cd44 | 2425 | 250 | 467 | 12 | 1430 | 1084 |
| Sema3a_Nrp2 | 4543 | 239 | 346 | 3 | 1009 | 562 |
| Ccn2_Itgam | 3315 | 209 | 447 | 7 | 1045 | 866 |
| Col1a1_Cd44 | 1427 | 208 | 378 | 12 | 1208 | 1086 |

Atm null with irradiation

| | n_spots | n_spots_sig | n_spots_sig_pval | n_cci_sig_celltype | n-spot_cci_celltype | n-spot_cci_sig_celltype |
|---|---|---|---|---|---|---|
| Fn1_Cd44 | 3572 | 276 | 663 | 8 | 2006 | 1521 |
| Col1a2_Cd93 | 1807 | 139 | 298 | 6 | 852 | 763 |
| Col1a2_Cd44 | 2432 | 133 | 276 | 7 | 887 | 678 |
| Ccn2_Itgam | 2142 | 110 | 232 | 6 | 575 | 465 |
| Spp1_Cd44 | 2846 | 101 | 302 | 7 | 671 | 433 |
| Nts_Ntsr2 | 1062 | 68 | 213 | 10 | 311 | 195 |
| Col1a1_Cd93 | 671 | 64 | 164 | 6 | 286 | 255 |
| Col1a1_Cd44 | 832 | 47 | 139 | 5 | 266 | 148 |
| Sema3a_Nrp2 | 1694 | 42 | 144 | 4 | 191 | 89 |
